# Computational neural mechanisms of goal-directed planning and problem solving

**DOI:** 10.1101/779306

**Authors:** Noah Zarr, Joshua W. Brown

## Abstract

The question of how animals and humans can solve arbitrary problems and achieve arbitrary goals remains open. Model-based and model-free reinforcement learning methods have addressed these problems, but they generally lack the ability to flexibly reassign reward value to various states as the reward structure of the environment changes. Research on cognitive control has generally focused on inhibition, rule-guided behavior, and performance monitoring, with relatively less focus on goal representations. From the engineering literature, control theory suggests a solution in that an animal can be seen as trying to minimize the difference between the actual and desired states of the world, and the Dijkstra algorithm further suggests a conceptual framework for moving a system toward a goal state. He we present a purely localist neural network model that can autonomously learn the structure of an environment and then achieve any arbitrary goal state in a changing environment without re-learning reward values. The model clarifies a number of issues inherent in biological constraints on such a system, including the essential role of oscillations in learning and performance. We demonstrate that the model can efficiently learn to solve arbitrary problems, including for example the Tower of Hanoi problem.

## Introduction

Consider a rodent making its way to a food source as a goal. The animal must remember where the food is, then plan a path to the food, and then execute a sequence of actions aimed at getting the food, all while ignoring distractors, avoiding dangers such as predators, and managing shifting food locations.

It is currently unclear exactly how the brain carries out goal directed behavior. Multiple neural mechanisms may drive goal-directed actions, including those of habits and more deliberate actions. Habits and reflexes offer a surprising amount of power for producing successful behavior, they can be inflexible and too tightly driven by the environment, and so cognitive control over behavior is necessary, as distinct from purely stimulus-driven actions. More developed nervous systems are capable of cognitive control, i.e. maintaining internal representations that can be used to guide behavior, partially decoupling the system from the external world. Goal-directed behavior is action driven by such internal representations of goals. William James (James, 1890, vol 2, pg 522) describes how actions must be driven sometimes by both a “movement’s sensible effects” and a “volitional mandate” which together drive the action. Later work cast a volitional mandate in terms of a rule that guides behavioral strategy in a given context (Miller & Cohen, 2001).

Cognitive control requires internal representations to drive behavior, which may include rules (Miller & Cohen, 2001), inhibition (Aron, 2007), performance monitoring (Alexander & Brown, 2011), or goal representation. The common theme is that cognitive control allows certain actions to be made while overriding otherwise salient cues that would drive different behavior (Stroop, 1935). While rules and inhibition may be useful in guiding behavior, they are not the same things as goals. Inhibition prevents inappropriate stimulus-driven actions. Rules define the ways by which stimuli are mapped onto particular actions, but rules are agnostic about end goals. By contrast, end goals define a desired state of affairs but can be agnostic about the actions required to achieve the goals. Studies of cognitive control have long focused on rules (Miller & Cohen, 2001) as well as inhibition (Aron, 2007) and performance monitoring (Alexander & Brown, 2011), but relatively little attention has been paid to how goals are represented and used to guide complex behavior.

Here we address how goals might be represented and drive behavior. The Goal-Oriented Learning and Sequential Action (GOLSA) model illustrates one way a neural system could plausibly learn to navigate an environment in pursuit of some goal. The network is comprised of continuous-time dynamic model neurons that utilize associative learning to construct a neural representation of an environment’s state space and use a variant of gradient climbing to reach a distant goal state from any starting state. Using principles from control theory, the GOLSA model address several key shortcomings of alternative accounts, including a unitary conception of reward and biological implausibility.

## Control Theory

Control theory describes how a system can regulate itself (Lee & Markus, 1967). At its simplest, it involves the negative feedback regulation of some variable, usually called a sensor signal, to maintain a particular set point value. A control system is constructed such that when the signal diverges from the set point, the system engages some effector to bring the signal back to the set point, where it should remain stable in the absence of external perturbations (also known as disturbances), despite inherent stochastic fluctuations. In practice, of course, perturbations are guaranteed, so the best case scenario for an effective control system is one that reacts to them quickly, accurately, and stably. Examples abound, such as the common thermostat which maintains a desired room temperature despite variations in the outside temperature, or the system by which a toilet tank maintains the desired full water level of the tank despite flushes, or short latency motor neuron reflexes which compensate for force perturbations.

A closely related concept is closed loop control, in which behavior creates sensory effects that drive subsequent actions (Powers, 1973). The fundamental insight of closed loop control is that behavior drives perception just as much as perception drives behavior. Agents behave so as to bring specific perceptual signals into the desired ranges (i.e., set points), such as the postural signals indicating a reach to a target location or the visual signals indicating that the agent has reached a desired location (Gibson, 1977; Hommel, Müsseler, Aschersleben, & Prinz, 2001). The concept has also been extended into value-based actions (Juechems & Summerfield, 2019), and is closely related to predictive coding (Friston, 2010) – at least in terms of minimizing the error between an actual and desired outcome, if not minimizing the error between the actual and predicted outcome (Ito, Stuphorn, Brown, & Schall, 2003).

Control systems generally have a limited range. Control theory offers a flexible and general account of goal-directed behavior, unifying the processes involved with a wide variety of natural and artificial systems. However, control theory assumes that a system is already in place to translate set point deviations into appropriate actions. While these systems are capable of flexibly adapting to changes in the range they were designed for, they are not capable of adapting their behavior in the face of unexpected changes.

In contrast, one of the greatest advantages afforded by a highly elaborated central nervous system is the remarkable adaptability to novel environments. When placed in a new spatial, social, or cognitive environment, an agent will typically not be able to achieve its goals. While the similarities between different environments of the same type will afford the agent some competence via generalization, navigation of the new environment will be difficult. With experience, performance will improve and the agent will be able to efficiently achieve its goals, conceived of here as bringing its perception into the ranges established by its goals, as predicted percepts (c.f. Friston, 2010). Additional mechanisms are required to model how this learning occurs, and here we propose how goal-directed behavior can be understood as driven by a more general, high dimensional control system, as inspired by control theory.

## Reinforcement Learning

Before describing the GOLSA model, we note that both model-based reinforcement learning (MBRL) and model-free reinforcement learning (MFRL) are ultimately inadequate to solve the general control theory problem. The basic issue is that with reinforcement learning (RL), goals are inflexible. There is typically one value assigned to a given state according to its proximity to reward. If the goal changes (i.e., the set point changes), then the reward values associated with a given state may no longer be accurate.

Both MBRL and MFRL suffer from this limitation. For example, Q-learning (Watkins & Dayan, 1992) associates a reward value (Q value) with each state and action pair. States closer to the rewarded (i.e. goal) state have progressively larger Q values. But what if the goal changes, and a different state is now rewarded instead? Then the Q values have to be relearned, which is inefficient. Alternatively, multiple Q values could be learned for each state, one per goal, but that would also be highly inefficient. At best, Q values might be approximated as a function of context (Wang et al., 2018). MBRL fares little better. To address more sophisticated cognitive abilities, MBRL agents use experience to learn an approximate model of the transition and reward probabilities in the environment and then separately learn a value (Q) function over the possible states of the world, using both value and model information to produce adaptive behavior. Like MFRL, MBRL still requires a learned Q value that cannot easily be re-learned as the rewarded state shifts.

One of the major limitations of all forms of RL is the way in which it construes reward, as essentially a scalar rather than a vector. Even Bayesian RL formulations which model the distribution of rewards still model the reward as the distribution of a scalar. This elides the distinct reward values associated with different goals. In the actual world, animals and people can learn based on receiving an unexpected outcome, even if our affective valuation of the alternative reward is identical to the expected one (McDannald, Lucantonio, Burke, Niv, & Schoenbaum, 2011).

Both of these issues can be addressed within the RL framework by redefining the states to include any relevant features such as reward identity, the level of various drives, etc. Because of the restriction that the reward signal must be a scalar, any complexity that *prima facie* ought to be included in affective signals instead must be offloaded to the more flexible state representations. However, this solution is not ideal. Practically speaking, it can result in a combinatorial explosion of states. It is also in tension both with the plurality of competing goods at the psychological level and the observation of multiple value signals at the neural level (Camille, Tsuchida, & Fellows, 2011; Kahnt, Heinzle, Park, & Haynes, 2011; Rudebeck et al., 2008; Rushworth & Behrens, 2008; Sescousse, Redouté, & Dreher, 2010). Even in the case of dopamine, the theoretical TD error signal may be a serious oversimplification of the midbrain dopamine system, which may represent more than one quantity (Dayan & Niv, 2008; Morris, Nevet, Arkadir, Vaadia, & Bergman, 2006; Roesch, Calu, & Schoenbaum, 2007).

## The GOLSA model

The Goal-Oriented Learning and Sequential Action (GOLSA) model (Figure 1) provides a novel account of goal-directed behavior that can handle many of the same tasks as reinforcement learning, using basic principles of control theory, Hebbian learning, and neural oscillations. Unlike in reinforcement learning, a GOLSA agent can select any known state as a goal to pursue rather than “reward.” A state is defined here as a particular set of conditions of the agent and its environment, which can be operationalized as a unique vector. States are represented by corresponding neurons in the GOLSA model. The goal state can be imposed externally or selected based on drive-reduction principles (Hull, 1943). Once selected, a goal-oriented map of the environment establishes a gradient of activity with a peak at the goal location (Ivey, Bullock, & Grossberg, 2011; Martinet, Sheynikhovich, Benchenane, & Arleo, 2011). The network then selects an immediate subgoal by selecting the state adjacent to the current state that brings the agent closest to the peak of the goal gradient. Once a desired next state is selected, an action is chosen to move the agent to the desired next state. This process is repeated until the goal state is reached. All learning in the model is essentially Hebbian, satisfying constraints of localist representation and broader neural plausibility. The model is conceptually similar to the A* (Hart, Nilsson, & Raphael, 1968) and Dijkstra (Dijkstra, 1959) algorithms commonly used in consumer GPS navigation devices, although the GOLSA model differs in that it respects neurobiological constraints (Supplementary Material).

**Figure 1.**
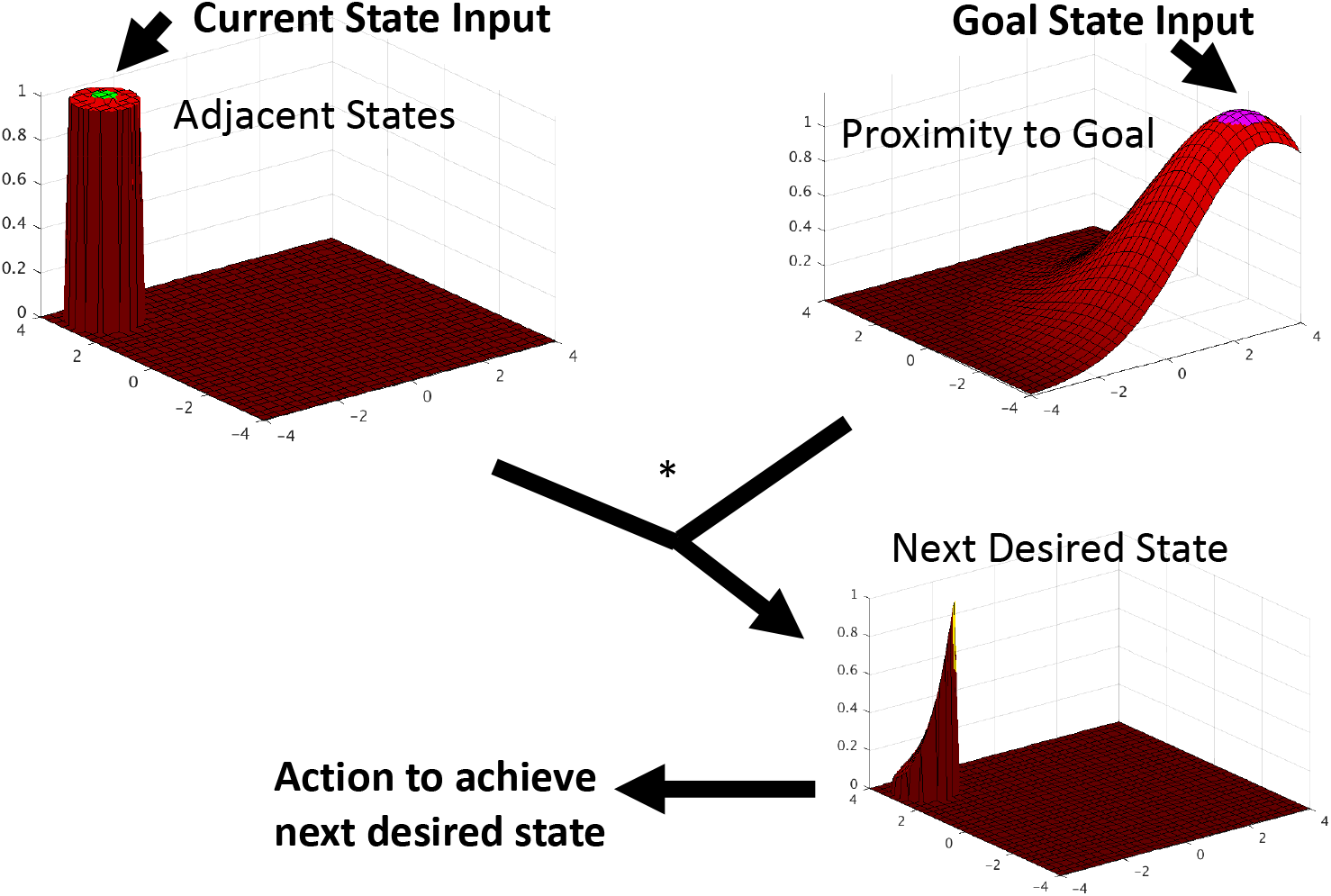
GOLSA model intuition. The model takes as input the current state and goal states and generates as output an action to move closer to the goal state. The model represents each state with a corresponding neuron in a given layer, and neurons representing nearby states that can be reached with a single action also share a synaptic connection. All state layers have representations of all possible states. The Proximity to Goal layer diffuses activity backward from the activated goal state. The Current State input activates all Adjacent States unit representations that are one step away from the current state. The Next Desired State is simply the unit-wise product of activities of the Adjacent States and Proximity to Goal layers, so that the most active unit represents the state that can be reach in one step and is closest to the goal state. Other model components (not shown) map the Next Desired state to an action aimed at achieving that state.

The GOLSA model has several distinct advantages compared to RL in the context of flexible goal pursuit. Chiefly, in RL a separate reward function must be learned for each goal, even if the layout of the environment is already known. The GOLSA model can take any possible state of the environment as a specified goal and plan actions to achieve it, without relearning the reward (Q value) function as would be necessary with RL.

The new model also provides a more concrete account of the neural plasticity required in goal pursuit tasks, which is underspecified in RL since algorithmic implementations offer the engineer substantially more flexibility than is afforded by neural implementations. Arrays of values in a computer system can be updated at any time using any other information in memory. In contrast, important values in neural systems necessarily have a biological substrate such as the weight of a synapse. There is no known (or plausible) mechanism for modification of arbitrary synapses based on information stored in distant structures or patterns of activity. Therefore, all modifications of synapses (or neural activity) must be due to information stored in a substrate physically present at that location. Though relatively straightforward, this constraint poses serious problems for neural instantiations of RL and other learning algorithms. The GOLSA model illustrates one way a neural system may operate to overcome this constraint in order to achieve flexible goal directed action.

## Methods

Figure 2A below shows the overall architecture of the model. Each red box in Figure 2B represents a layer of rate-coded artificial neurons which together serve a particular function of the model. Depending on the layer’s function, each unit represents either an environmental state, an action the agent can take, or a state transition. The grey ovals represent nodes which provide external input or control signals to the model that are not represented in terms of neural activity. Many of the projections between model layers are hard-coded one-to-one mappings, though several key projections are fully plastic and instantiate the learning processes described in detail below.

**Figure 2.**
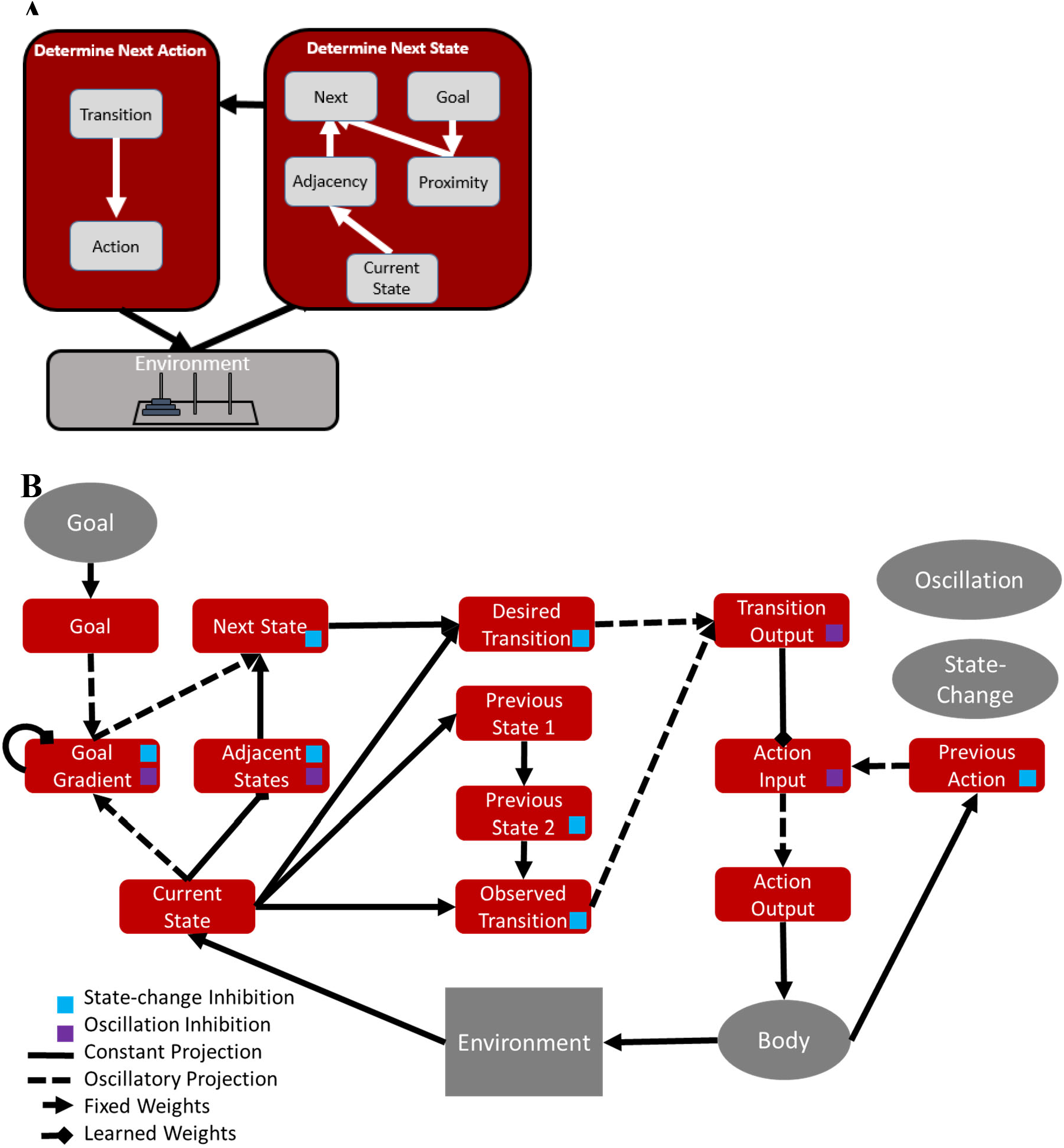
A) Schematic overview. The right box encapsulates Figure 1 above, while the left box depicts the desired transition (conjunction of current state and desired next state), which activates a corresponding action (i.e. an inverse model). This in turn causes changes to the current state of the agent in the environment. B) Full diagram of core model. Each rectangle represents a layer and each arrow a neural projection. The body is a node, and two additional nodes are not shown which provide inhibition at each state-change and oscillatory control (below and Supplementary Material). The colored squares indicate which layers receive inhibition from these nodes. Some recurrent connections not shown.

## GOLSA Algorithm

Though the GOLSA model is comprised primarily of continuous time dynamical model neurons and synapses, its behavior can be roughly described by the following algorithm.

1. Determine which states can be reached from the current state.
2. Identify one of these states that will best bring the agent closest to the current goal.
3. Take the action that will likely implement the transition between the current state and the desired next state.

This process requires learning the adjacency structure, i.e., the topology, of the state space as well as a mapping from transitions to actions. This information is instantiated in the connection strengths of three key projections between layers in the network: from *current-state* to –*adjacent states*, from *goal-gradient* to itself, and from *transition-output* to *action-input*. A description of model activity after successful learning is presented below, followed by an explanation of the learning process. For ease of visualization, the model activity will generally be described in the context of a six-state gridworld arranged as shown in Figure 3, where each state can be reached from the states physically adjacent to it. Although this appears identical to a simple 2D navigation problem, the approach generalizes to high dimensional states representing arbitrarily complex problems. A more complete description of the model is in the Supplementary Material.

**Figure 3.**
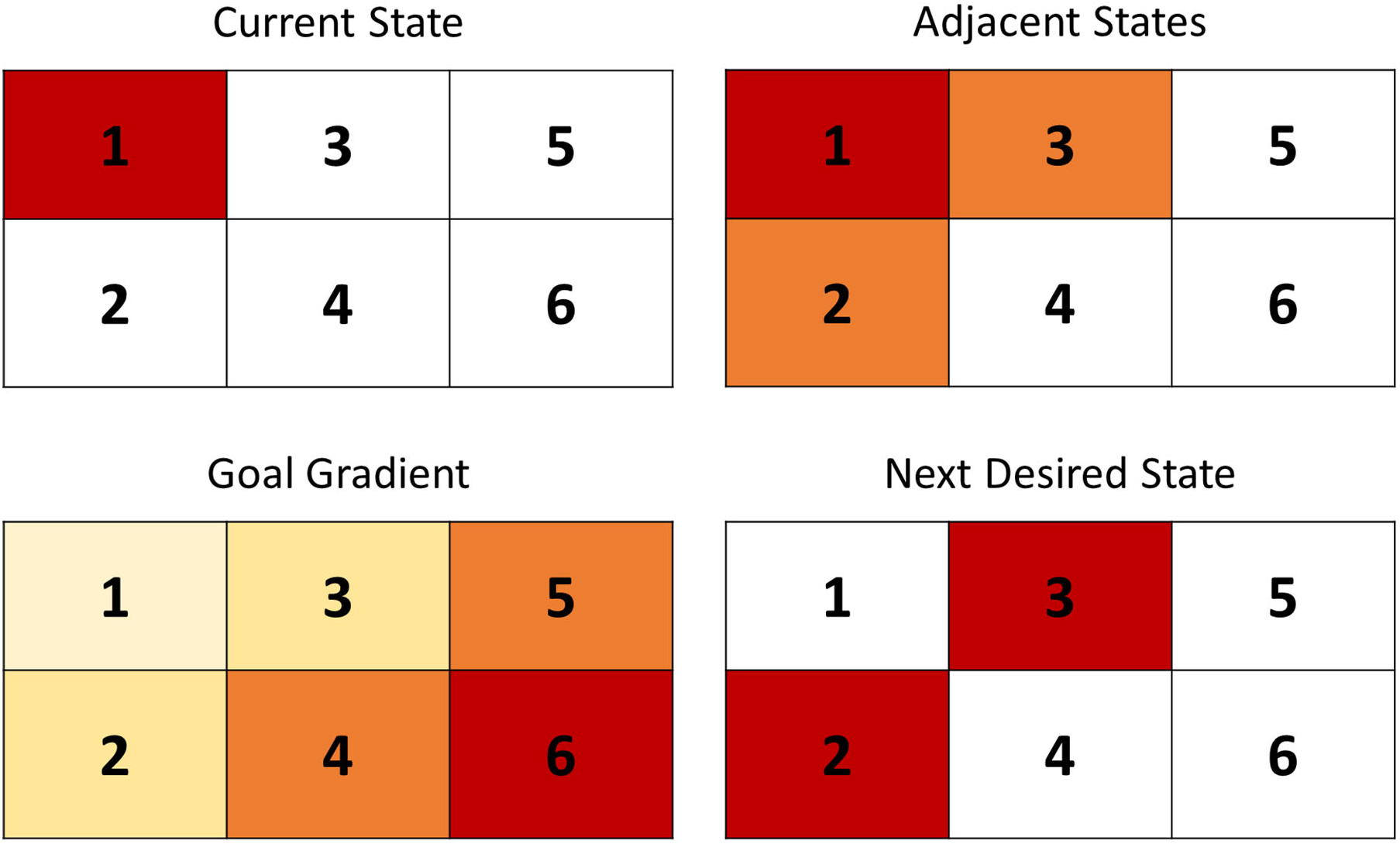
GOLSA model example and Illustration of important model layer activities for selecting the desired next state. Each box corresponds to a unit in the appropriate layer and more red units are more active. In the current-state layer, only the unit corresponding to the agent’s state is active. In the adjacent states layer, units representing states one step away are also active, but less so. Each unit in goal-gradient in rough proportion to the proximity to the goal state. When these two maps are integrated in next-desired-state, the state(s) which bring the agent one-step closer to the goal are activated.

Figure 3 illustrates the basic mechanisms of the algorithm with a simple example. Each of the four maps of the environment corresponds to one of the layers in the network, and each of the six boxes in the map represents the corresponding unit in the layer. Each layer represents the same six states. The colors indicate the level of activity expected in each unit for a trial in which the agent starts in state 1 and has the goal of navigating to state 6.

For the representation of the current state, only the unit corresponding to the agent’s actual location should be active (Figure 3). Naturally, the representation of adjacent states includes all and only the states reachable from the current state, including the current state itself. Within the *adjacent-*states layer, the current state is actually more active than the adjacent states to facilitate learning (Supplementary Materials). The goal gradient has a peak of activity at the goal location (state 6) and then a smoothly decreasing level of activity diffusing outward from the goal. This representation encodes each state’s distance from the goal as a pattern of activity. The goal gradient and adjacent state representations interact to produce a representation of the next desired state. The pattern of activity in this layer answers the question, “Of all the states adjacent to the current state, which shows the most activity in the goal gradient?” Here there are two states that are both adjacent to the current state and equally close to the goal state. In the actual network, a winner-take-all (WTA) system breaks the symmetry and selects a single state to move to next. Using the current state and the next desired state, the network identifies the appropriate transition to implement and then takes the appropriate action based on a learned association between actions and transitions. If the goal gradient is thought of as a three-dimensional hill, the model climbs the hill by successively enacting the transitions that move the agent as far up the hill, and toward the goal, as possible.

## Learning

The core model, described above, depends upon accurate learning in three projections, namely from *current-state* to - *adjacent-states*, from *goal-gradient* to itself, and from *transition-output* to *action-input*. (Figure 2B) All other connections utilize hard-coded mappings with a one-to-one or simple combinatorial structure.

During early learning, the agent lacks knowledge about the structure of the state space and will therefore be unable to determine an appropriate next desired state or the action required to take an appropriate transition. Without determining the correct transition, the agent will obviously fail to take the appropriate actions to reach the goal. In our implementation of the model, after 300 time steps without taking an action, the agent “explores” by taking the available transition that has been implemented least often. Figure 4 shows model activity over the course of the first transition in a trial prior to any learning with the same starting and goal states as in Figure 3, namely state 1 and state 6.

**Figure 4.**
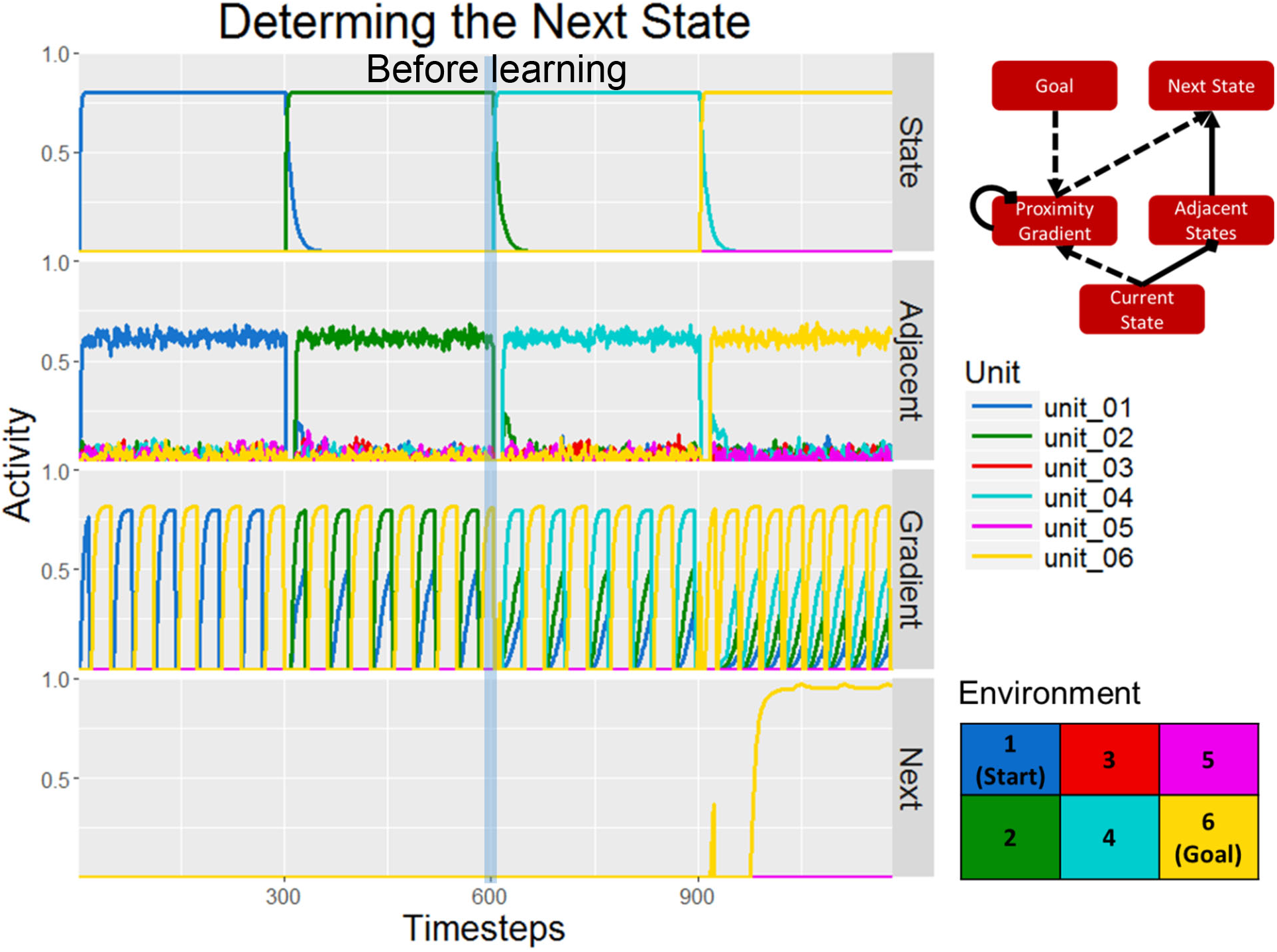
Time Steps 0-600: Layer activities related to determining the desired next state during the first step of the first trial during learning. As described in Figure 3, the model is starting at state 1 with the goal of state 6. No states are known to be adjacent so only current state activity is present there in the Adjacent layer. After the first transition, the goal gradient learns as an oscillatory signal (describe below) provides input from the current State, which is now state 2. The model thus learns (in connections within the goal gradient layer) that state 1 can precede state 2. **All time steps 0-1200:** Network activity related to determining the next state during the entire first trial, before learning has occurred. The trial continues until the agent reaches the goal state via random exploration. Note that when the agent reaches state 6, the goal, the entire trajectory is encoded in the goal-gradient layer. All previously visited states are active in order of their proximity to state 6.

As shown in Figure 4 (time steps 0-600), the *adjacent states* layer initially mirrors the *current-state* layer, while the *goal-gradient* alternates between a simple representation of the current state and the current goal. Without the necessary information to determine the appropriate next state, the agent explores, moving down into state 2. This results in learning in both the projection from *current-state* (the State 1 unit) to *adjacent-states* (State 2 unit) and the recurrent projection in *goal-gradient* (from State 2 unit to State 1 unit). Only the latter has a visible effect on the activities here because the *goal-gradient* represents backward transitions while the *adjacent-states* layer looks forward into the future. Because the model does not assume that all transitions are bidirectional, it does not yet know that it is possible to move from state 2 to state 1. The *goal-gradient*, on the other hand, now contains unit 2 activity during the learning phase, indicating that the model (specifically the *goal-gradient* layer) now knows that if it needs to get to state 2, it can do so via state 1.

The activity across the full first trial, shown in Figure 4, demonstrates how the gradient continues to expand with experience. Because the agent happened upon the goal state, the trial ended. Note that until the agent actually reaches the goal state, no unit is active in *next-desired-state* and therefore no *desired-transition* is specified. This is because during most of the trial, no units were active in the *goal-gradient* during the action-phase of the oscillation at the same time that the corresponding unit was active in *adjacent-states*. Once the agent reaches state 6, this is no longer true and state 6 becomes the desired state and the 6-to-6 transition unit is activated.

For most of this trial, no desired next state was specified. Therefore, the network could not select a desired transition to implement with an action. This is shown in Figure 5 with the lack of activity in *desired-transition* and *action-output* for most of the trial. Recall that transition layers dedicate one unit to each combination of starting and next states, such that there are 36 units in each transition layer for this simulation. Unit 1 represents the transition from state 1 to state 1 while unit 36 represents the transition from state 6 to state 6. Unit 36 becomes active at the end of the trial, when a unit finally becomes active in *next-desired-state* and the agent both currently inhabits state 6 and desires to be in state 6 at the next step.

**Figure 51.**
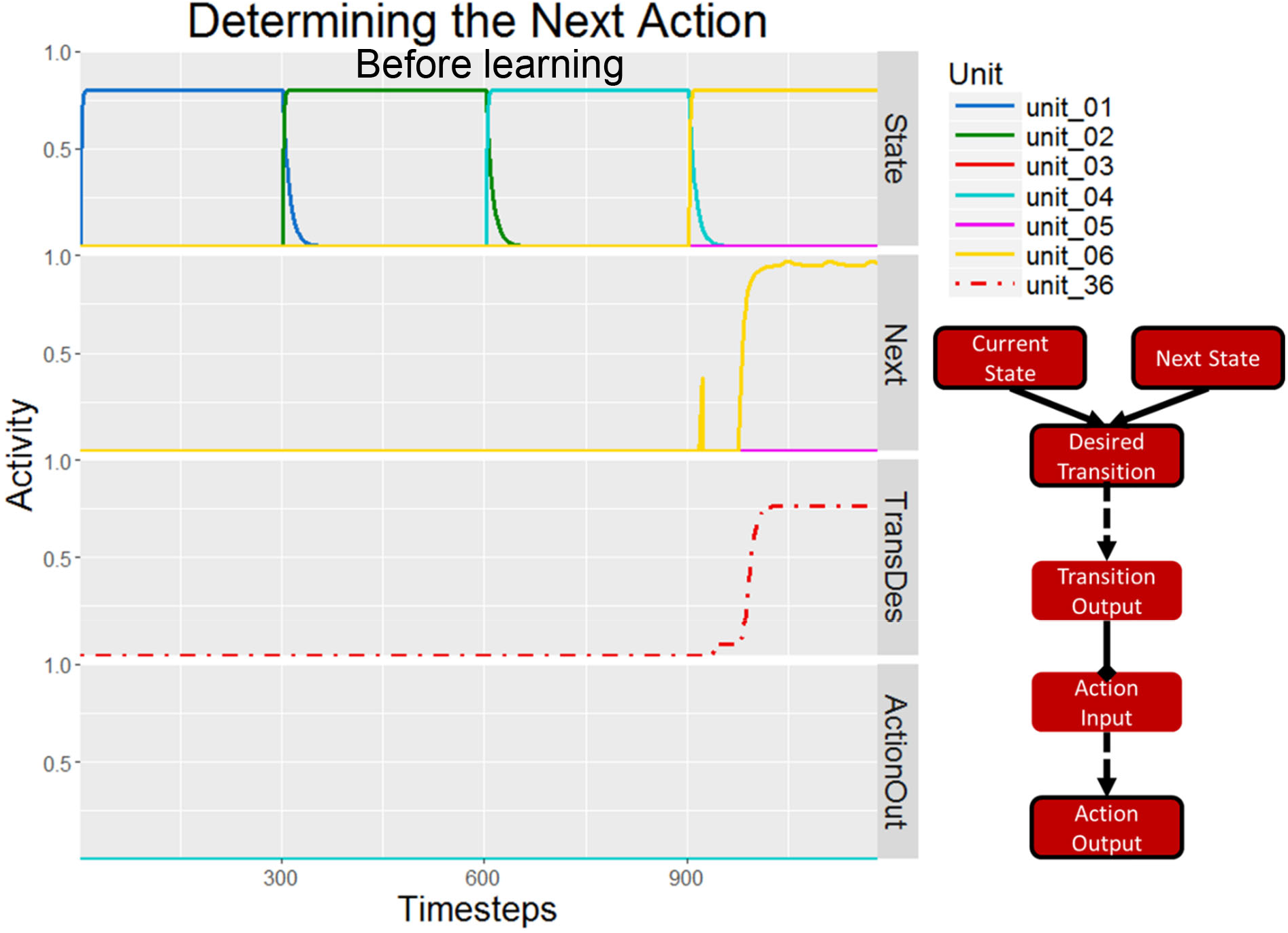
Network activity in layers related to action selection during the first trial during learning, before learning has occurred. Note that transitions are numbered such that transition 1 is from state 1 to state 1, transition 6 is from state 1 to state 6, transition 7 is from state 2 to state 1, etc. Because no next state is specified during most of the trial, no transition is specified and no action is activated.

On the next trial (Figures 6, 7), the agent is able to use the updated weights to walk the same path to the goal more quickly and without using forced exploration.

**Figure 6.**
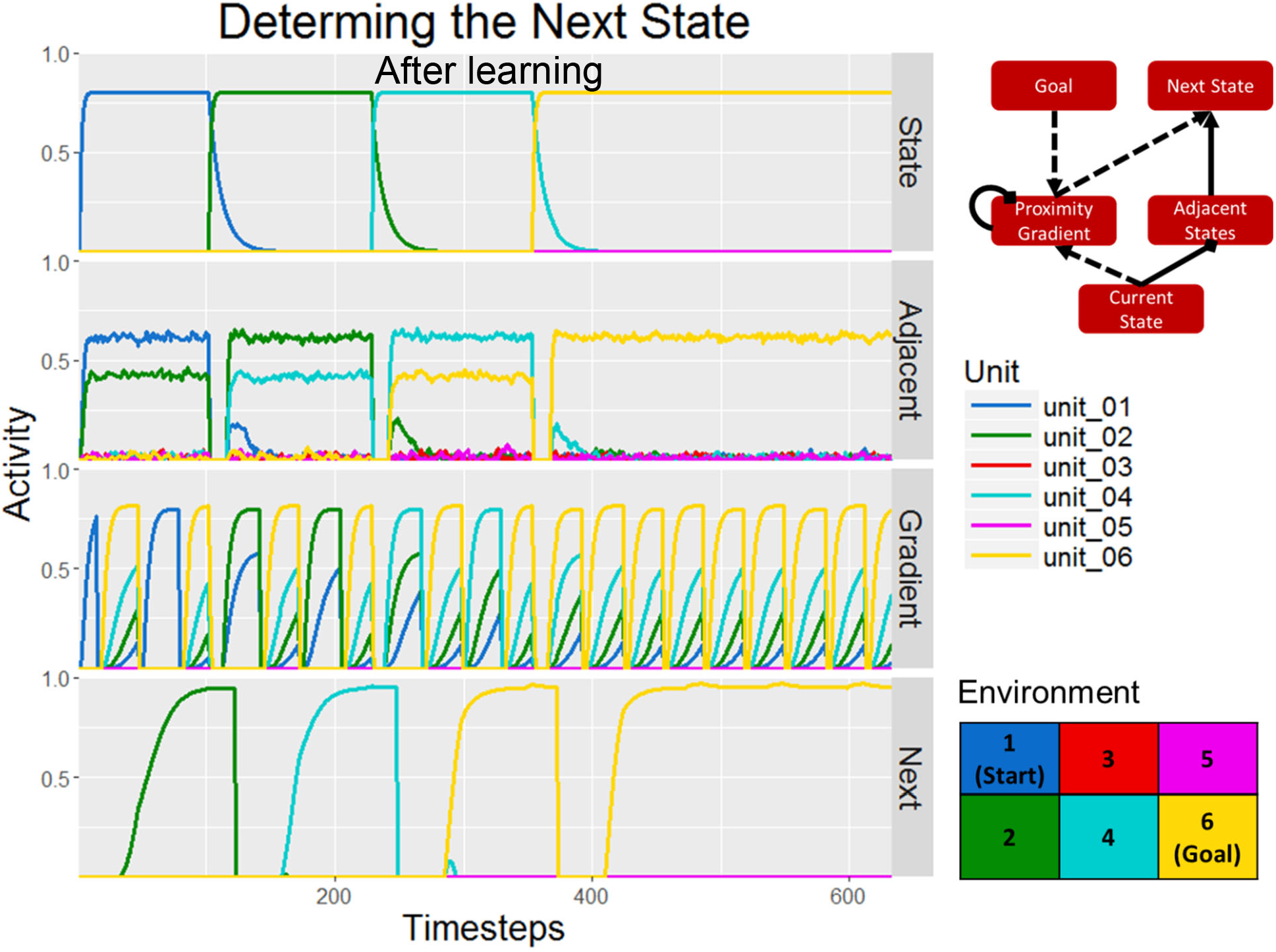
Network activity in the layers responsible for goal selection during the second trial after learning. The agent utilizes the past experience in the environment to navigate back to the goal path. In each state, the appropriate next state is determined based on the activities in the adjacent-states layer and the goal-gradient layer. Gradient activity is only used during an oscillatory phase in which the goal rather than the current state is an input to the gradient.

**Figure 7.**
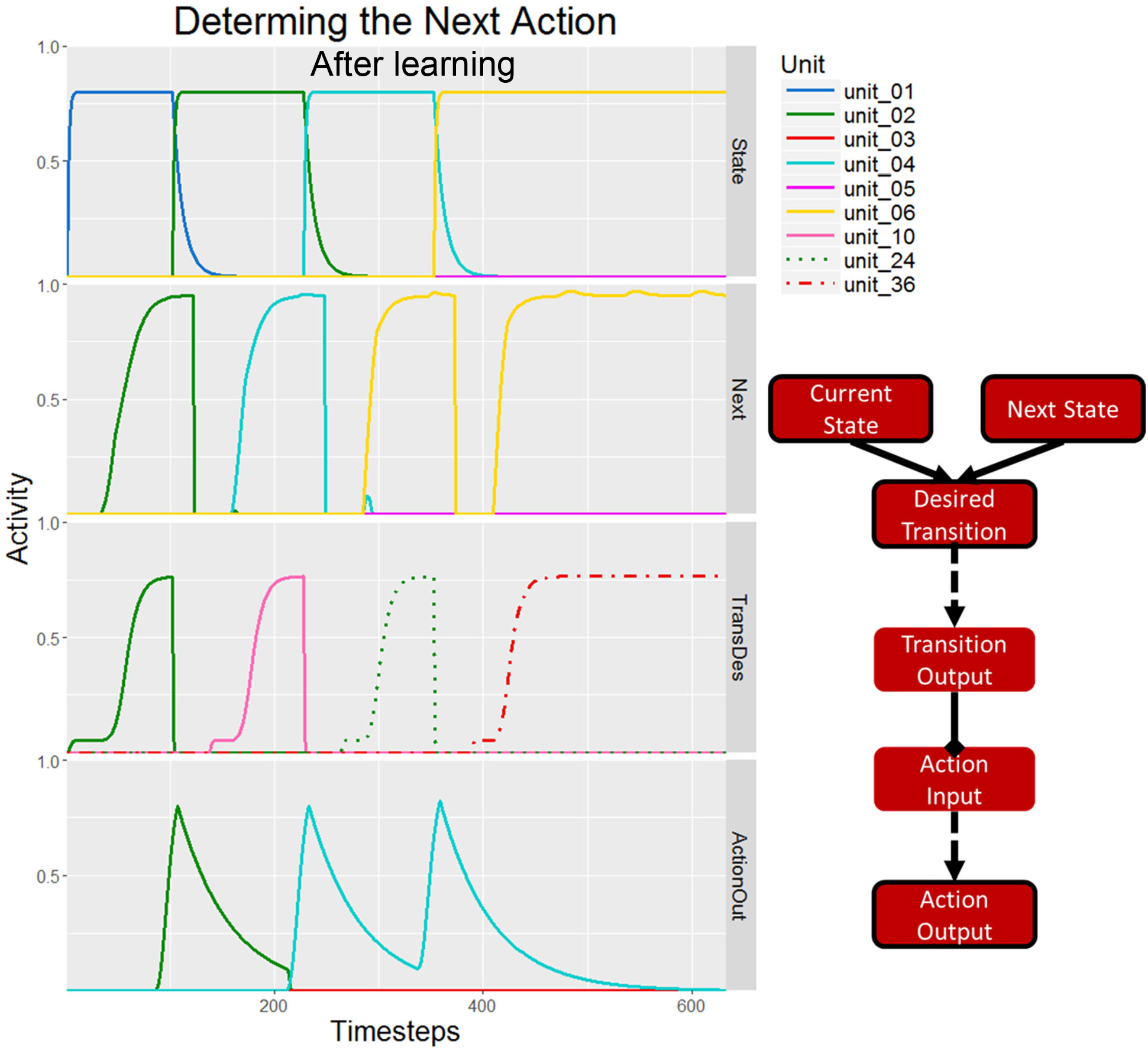
Network activity in layers related to action selection during the first trial during learning. In the first trial, shown in Figure 51 5, a desired transition was not specified for most of the trial because a next-desired-state was not selected. In this trial, the specification of a next-desired state (Figure 6), results in the specification of a transition in desired-transition (TransDes), which drives the appropriate actions compared to pre-learning where no actions were selected (Figure 5).

**Figure 8.**
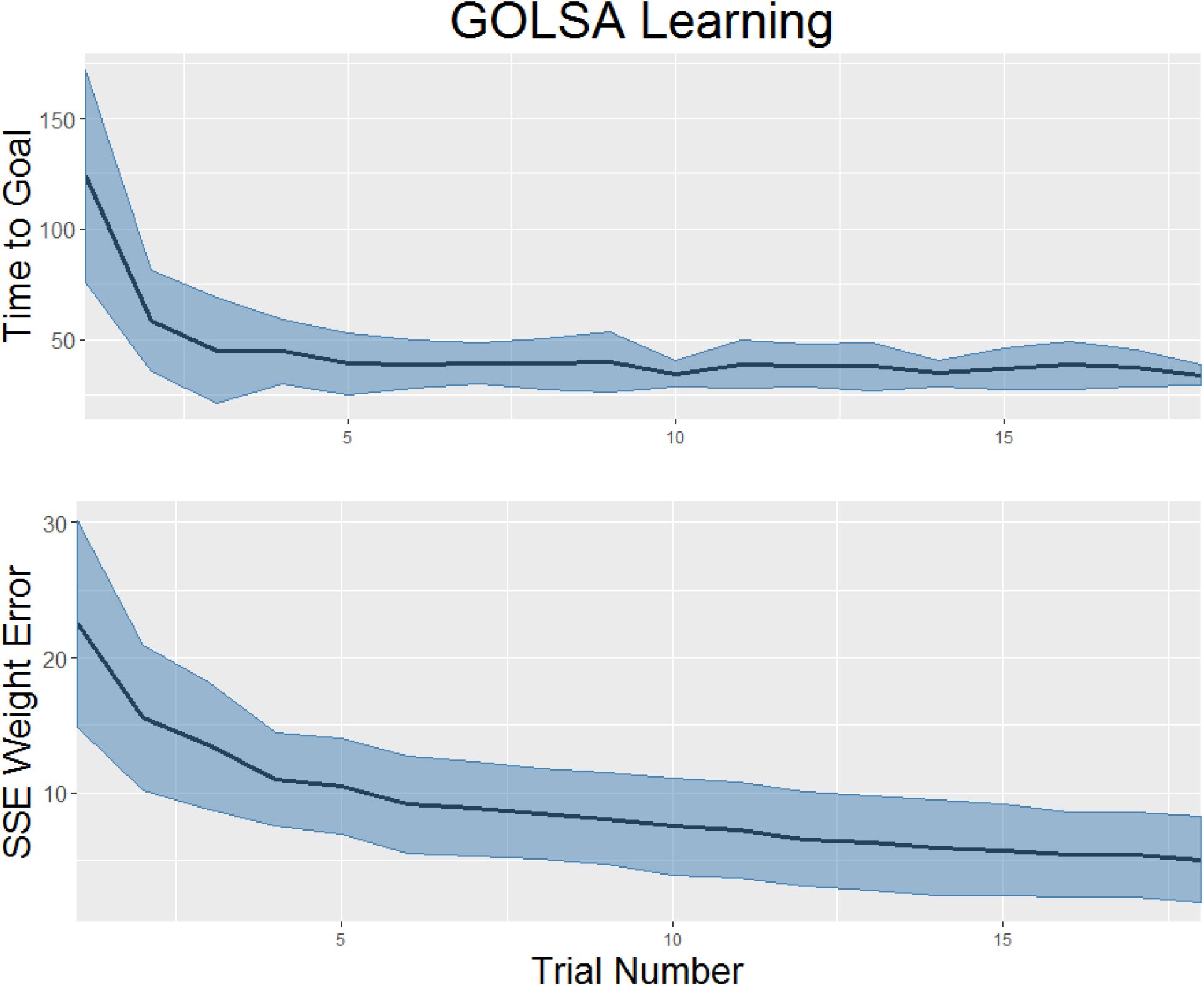
**Top:** Average time taken to the goal as a function of the number of trials since learning began, in the six-state gridworld problem. The learning process was repeated 50 times and the shaded area represents the standard deviation across those 50 trial sequences. **Bottom:** Total sum squared error of the learned weights (actual – ideal) in the three key projections described above.

On this trial, the *goal-gradient* and *adjacent-states* layers contain enough information to activate the correct unit in *next-desired-state* and therefore in *next-desired-transition*. The agent’s exploration in the previous trial taught it which actions implement those transitions, so activity is present in *action-input* during the acting phase (the smaller peaks) in addition to the previous action information during the learning phase (the larger peaks). This activity accumulates in *action-output*, which eventually triggers the *body* node to effect an action. The following section addresses the learning laws which allow the agent to move from blindly exploring in first trial to effectively navigating to the goal in the second trial.

## Oscillations

The constraint of biological plausibility leads to a problem with learning the mapping from desired state transitions to actions. Specifically, the *dual roles problem* requires the same projection, e.g. the mapping from desired state transitions to actions, to be presented with different information at different times, within a single event, for learning vs. performance. Consider that when a desired transition node is active, indicating a desired transition from, say, state 1 to state 3, the action (A) that is generated may instead cause a transition from state 1 to state 2. In that case, it would be inappropriate to strengthen the active synaptic weight from a node representing the desired transition (1➔3) to the action A, based purely on a Hebbian principle of coactivation. Instead, we would like to activate a node representing the actual transition (1➔2), and then strengthen its projection to the actual action A. This leads to another problem, which is that if we activate a representation of the desired transition (1➔ 2), what is to prevent that from activating some other action inappropriately? All of this suggests that the network must operate in two distinct modes, i.e. learning vs. acting. During the acting mode, the desired state transition must activate an action, which will then be generated. During the learning mode however, two changes must be made. First, the activity pattern of the desired transition layer must be changed to represent the *actual* transition that just occurred, instead of the *desired* transition. Second, the action generating layer must have its output disconnected, so that it cannot generate an inappropriate action during learning. To fulfill this requirement, we posit an oscillatory mechanism that rapidly switches the network between the learning and acting modes described above, with cell activities reliably linked to an oscillatory cycle (Klausberger & Somogyi, 2008), reminiscent of a computer’s CPU clock cycle. The details of this mechanism, and its application to the three learned connections in the model, are described in the Supplementary materials.

## Results

### Small Environment

Over the course of multiple trials, the GOLSA model learns how to accurately navigate its environment to reach any goal state from an arbitrary starting state. Notably, it can generalize, learning to reach goal states without having been on those states as goals. With a constant goal and starting state across trials however, the model would simply repeat the first path it found via random exploration. To counter this tendency, the model was run with alternating starting and goal states. On each trial, it attempted to get from one corner of the small 6 state environment to the other, forcing the agent to take different paths to reach its goals. The agent also had a 20% chance to randomly take a relatively unexplored transition when attempting to take a known state transition. A trial was ended shortly after the goal was reached, or after 4000 time steps, with each time step equal to 50 msec (t = 200 seconds total) if it was not. Figure below shows how the average time to complete a trial and the overall error in the learned projections decreased over the course of 18 trials. The entire sequence of trials was repeated 50 times.

As expected, the time taken in each trial diminished rapidly to a minimum of about 30 time steps, though random exploration sometimes moved the agent off course making each trial take longer. The total sum squared error in the learned weight projections also diminished rapidly over the course of the trials, though failed to completely reach zero because some transitions were never explored.

### Tower of Hanoi

The GOLSA model can learn to navigate small gridworlds, but can it learn to navigate larger, real-world problems? To test this, we taught the model to complete the Tower of Hanoi puzzle. The Tower of Hanoi consists of three differently-sized discs on three pegs. The typical starting configuration is all three discs on the leftmost peg. For a position to be legal, smaller discs cannot be below bigger discs. On each step, participants move the top disc from one peg to another, with the typical goal state being all three discs stacked on the rightmost peg. There are 27 legal states in the three-disc Tower of Hanoi. These states can be arranged into a graph where each node constitutes a legal state and each link represents a valid move.

To put the problem into a format that the GOLSA model can complete, each possible legal configuration was considered a state. There were 6 possible actions, each defined as attempting to move the top disc from one peg to another. The only time this “action” would fail is if there was in fact no disc on the “from” peg, in which case the action could still be taken but would not result in a state transition.

The architecture used was very similar to that used in the small six-state grid world, but with larger layers containing units to represent each of the 27 states or 729 transitions. The decay parameter on the *goal-gradient* was also changed from 0 to 1 (effectively turning off the decay), allowing goal-related activity to diffuse more broadly through the network. Without this change, the decay would overwhelm the relatively small effect of goal activity passing backward through many states in a trajectory. While the gradient could still be successfully established after exploration-based learning strengthened the majority of the appropriate weights, a single learned trajectory could not be reliably followed even if repeated many times. After removing the decay term (and adjusting the learning thresholds accordingly), however, the gradient is much more sensitive.

Learning proceeded in a similar fashion to that used in the small gridworld. While the three discs on the right peg constitute the typical goal state for the Tower of Hanoi, a major strength of the GOLSA model is its ability to navigate to any desired state. To facilitate learning over the entire space, we rotated the starting and goal states across three configurations during which the agent attempted to move from one corner of the state space show in **Error! Reference source not found.**A clockwise to the next corner. The max length of a trial was 200 time units (4000 timesteps at dt = .05), and the agent could remain stuck for a maximum of 15 time units before exploring. Though the environment was significantly larger than the six-state grid-world, the model quickly learned the topology of the state space and the actions required to traverse it, as illustrated in Figure 9B.

**Figure 9.**
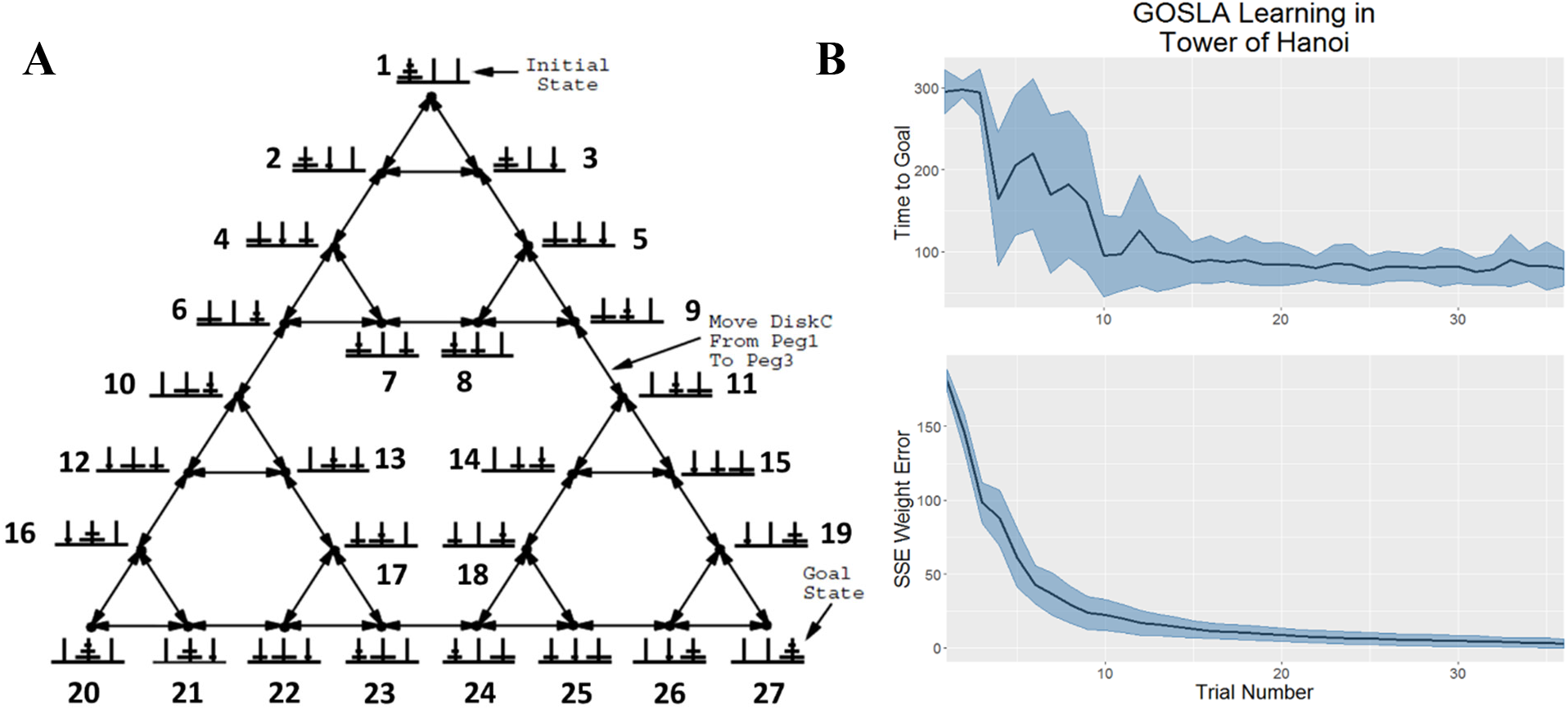
A) Graph representation of the Tower of Hanoi state-space. States are numbered according to their representation in the model (e.g. state 22 is represented by the 22^nd^ unit in current-state, adjacent-states, etc. Figure adapted from Knoblock (1990). **B)** Top: Average time taken to the goal as a function of the number of trials since learning began. The learning process was repeated 50 times and the shaded area represents the standard deviation across those 50 trial sequences. Bottom: Total sum squared error of the learned weights (actual – ideal) in the three key projections described above.

The GOLSA model demonstrates how goals can be effectively pursued after biologically plausible learning. However, it obviously has a much simpler architecture than a real biological cognitive system. Unsurprisingly, there are a few major ways in which performance on the task diverges from human performance. One challenge humans face on the Tower of Hanoi and other similar puzzles is that people can get stuck in situations where they appear to have to move away from the goal state in order to eventually reach it (Goel & Grafman, 1995). For instance, even though all discs must eventually end up on the third peg to reach the classic goal state, discs must be moved off the third peg several times. The problem becomes even more pronounced when more discs are added.

This difficulty is due to the fact that humans can represent the problem space in multiple formats. Each state has a visual representation in addition to a more abstract cognitive representation. However these representations diverge in that two states may be visually similar yet quite distant in the true state space. The visual representation may require fewer cognitive resources than an abstract representation of each state, potentially leading people to make inefficient moves. In contrast, the GOSLA model only represents the state space in terms of its transitions. It cannot be confused by a problem that requires one to “go back before going forward” as progressing toward the goal state is always represented as moving forward. Perhaps a more realistic, though less efficient model, would include two goal layers, one which represents the objective goal which provides input to a “visual-goal” layer such that states which are visually similar to the objective goal are activated.

## Discussion

The GOLSA model implements a hill-climbing approach to goal-directed behavior in which an agent ascends a gradient distributed across state-space with a peak at the goal. The model also strictly adheres to biological constraints including localist learning laws. To establish the gradient and learn which paths are available, the network learns the topology of the state space based on experience, using Hebbian learning modified by eligibility traces and oscillatory dynamics. Oscillations are necessary because the *dual roles problem* requires the same projection, e.g. the recurrent projection in *goal-gradient*, to be presented with information about the past (where it had been) during learning but actually perform the necessary computations on information about the future (where it is going). In the language of control theory, the goal serves as a set point for the network which then attempts to align its current state with that goal.

Additional mechanisms described in the Supplementary Material allow the model to learn, via standard reinforcement learning, to associate certain homeostatic drives with goal states that satisfy those drives. In this way, homeostatic drives can determine the set point of a control system. As actions bring the system to the goal state, the homeostatic drive shuts off. This effectively constitutes a negative feedback control loop in the sense of control theory. Reinforcement signals occur when a homeostatic drive is sated, so that drives which are shut off as reward is achieved in a goal state lead to a strengthening of the connection from the (sated) homeostatic drive to the goal state in which the drive is satisfied (Hull, 1943).

When the goal is specified externally, the model can seek any state in the space without any need of relearning. This is an advantage over Q-learning approaches in which values are intrinsically tied to corresponding states. The GOLSA model separates the value of a state from the cognitive map that defines what state transitions are probable, and both the value of a state and the probability of state transitions are further separated from action selection. The separation of values, states, and action representations allows them to be flexibly combined as needed, which affords the arbitrary goal specification ability (and consequent revaluation of states) that is generally lacking in both model-based and model-free RL approaches.

An additional competitive queueing module (Supplementary materials) allows the network to simulate a sequence of state transitions before executing any actions, and then store the appropriate transitions for future execution. The model is capable of successfully navigating simple environments, including abstract environments like the state space of the Tower of Hanoi, with relatively little learning.

### Oscillations

Neural oscillations play roles in a wide variety of cognitive functions including memory, cognitive control, sensorimotor integration, and behavioral timing (Berger, 1929; Caplan et al., 2003; Cavanagh & Frank, 2014; Coon et al., 2016; Herrmann, Munk, & Engel, 2004; Palva, Monto, Kulashekhar, & Palva, 2010; Roux & Uhlhaas, 2014). Oscillations play three major roles in the GOLSA model: gating of projections, gating of learning, and inhibition, as we describe below.

### Projections

A key part of the GOLSA model is the compression of learning and acting phases into distinct oscillatory phases of activity interleaved within a single behavioral run as opposed to separated between runs. The dual roles problem requires that different patterns of information pass through certain projections in the two distinct phases, and this is accomplished with projections that ‘open’ and ‘close’ based on the phase of an oscillatory node. An open projection passes the weighted presynaptic input to the postsynaptic layer while a closed projection prevents any input from passing through. A similar scheme may be instantiated in the brain via cross-frequency coupling, in which oscillations at different frequency bands interact to facilitate long-range communication (Ryan T. Canolty & Knight, 2010; Roux & Uhlhaas, 2014). Cross-frequency coupling typically involves an interaction between the phase or amplitude of one oscillation, and the amplitude, power, or frequency of a second oscillation (Jensen & Colgin, 2007). Cross frequency coupling has been directly observed in animals (Buzsáki et al., 2003; Hentschke, Perkins, Pearce, & Banks, 2007; Lakatos, 2005; A. B. L. Tort et al., 2008) as well as humans (Bruns & Eckhorn, 2004; R. T. Canolty et al., 2006; Cohen, Elger, & Fell, 2009; Mormann et al., 2005). This phenomenon can facilitate communication by providing a phasic window for input from particular cell types onto target neurons (Klausberger & Somogyi, 2008). Closing a gate in the model is therefore analogous to nudging the phase of the source or target population such that signals sent from the former to the latter would arrive outside the temporal window for effective communication.

### Learning

While our understanding of the relationship between neural oscillations and synaptic plasticity is still quite limited, a substantial body of work has focused on the trisynaptic pathway in the hippocampus between entorhinal cortex (EC) and hippocampal regions CA1 and CA3. Long term potentiation (LTP) in dentate gyrus and area CA1 of rats is modulated by the relationship between the timing of input and the phase of theta oscillations in those regions (Hölscher, Anwyl, & Rowan, 1997; Huerta & Lisman, 1993; Hyman, Wyble, Goyal, Rossi, & Hasselmo, 2003; Orr, Rao, Houston, McNaughton, & Barnes, 2001; Pavlides, Greenstein, Grudman, & Winson, 1988). This effect is reproduced in a simplified fashion in the GOLSA model, in that the recurrent connections within *goal-gradient* undergo synaptic plasticity only when the *substitution-oscillation* is in a particular phase. Hasselmo et al. (2002) provide a model of CA1, CA3 and EC using a very similar mechanism to illustrate how theta phase can control distinct phases of information encoding and retrieval. In addition, context learning in rats correlates with the strength of cross-frequency coupling in CA3, in particular the effect of theta phase on gamma amplitude (a. B. L. Tort, Komorowski, Manns, Kopell, & Eichenbaum, 2009).

### Inhibition

Intra-layer inhibition plays an important role in establishing WTA competitions in the GOLSA model. Layer-wide inhibition is also extremely important to model function, and serves a number of roles, such as resetting each layer for new input and facilitating new WTA winners. During online performance, inhibition is largely driven by the *state-change* node which attains a high value when a new state is reached and then decays, prompting layer-wide inhibition in different layers as its value drops. Layers are typically inhibited immediately before new information is gated in. It is crucially important that gating and inhibition for each layer occur within the proper phases of the oscillation. For instance, *next-desired-state* must be cleared via inhibition after *goal-gradient* and *adjacent-states* are cleared because *next-desired-state* depends on the accuracy of the patterns the other two layers and those patterns take time to establish themselves after inhibition.

Oscillations and inhibition are tightly coupled in the brain. Oscillations appear to be driven in large measure by the rhythmic dynamics of inhibitory interneurons while at the same time driving various inhibitory processes (Isaacson & Scanziani, 2011). On a large scale, alpha power as measured with EEG appears to drive inhibition of entire cortical regions, similar to layer-wide inhibition in the model (Jensen & Mazaheri, 2010; Klimesch, Sauseng, & Hanslmayr, 2007).

#### Reinforcement Learning

The GOLSA model is not a direct competitor to MBRL. As presented, the GOLSA model assumes that state transitions and rewards are deterministic rather than stochastic, and there is no unitary conception of reward as in an MDP. However, to the extent that MBRL attempts to explain how organisms pursue goals in their environment, it is closely related to the GOLSA model. Both approaches require learning similar information. The projection from *drives* to *goals* (Supplementary material) is analogous to the reward matrix in MBRL in that it specifies which states are likely to lead to rewards, conceived of here as a reduction in drives. They are different, however, in that drive reduction occurs deterministically and there is no limit to the number of possible drives, whereas, crucially, there is typically only one kind of reward in an RL context.

The MBRL transition matrix specifies a probability for each transition (s, a, s’). A synapse in a neural network inherently relates two quantities rather than three, typically the firing rates of the pre and postsynaptic neurons. As a result, the representation of transition probabilities in the model relies on three projections. The first two simply map combinations of current and desired next states to transitions. Each unit in *current*-*state* excites the units in *desired*-*transition* which represent transitions that begin at the current state. Likewise, the units in *next*-*state* excite the *desired-transition* units representing transitions which terminate at the desired next state. Together with a WTA as a result of lateral inhibition within *desired*- *transition*, these hardcoded projections result in the unique specification of transitions (s, s’). *Desired*-*transition* connects one-to-one with *transition*-*output* which in turn projects to *motor*-*input* via a learned projection. The learning goal for the synapses in this projection is to indicate which transitions ((s, s’), a) are viable.

Action selection in the GOLSA model also proceeds in a substantially different fashion than in RL in that the GOLSA model does not explicitly assign values to actions in states. Instead, action is driven first by the selection of a desired next state, based on which available next state will take the agent closer to the desired goal. From there, a desired transition is specified and the action which will implement this transition is taken.

The major differences in action selection between MBRL and GOLSA necessitate different uses of learned information about the environment. In MBRL, the transition and reward matrices are most commonly used to perform forward sweeps through possible action plans (Moore & Atkeson, 1993; Peng & Williams, 1993; Sutton, 1991). The simulated reward (or lack thereof) on each sweep can then be used to update the state or action values. This process allows for greater flexibility compared to model-free methods since, for instance, the agent can immediately devalue states preceding a newly discovered roadblock rather than waiting to actually enter those preceding states on a future trial.

In contrast, the GOLSA model does not use forward sweeps to aid learning. Instead, information about the structure of the environment is used directly to determine the desired next state. The information used in this procedure is not the transition-to-action mapping in the projection from *transition-output* to *action-input*, analogous to the learned transition matrix in MBRL. Instead, the desired next state is selected using the state adjacency information instantiated in the weights of the projection from *current-state* to *adjacent-states* and the recurrent connection from *goal-gradient* to itself. The former represents the forward-time adjacency structure whereas the latter represents the backward-time adjacency structure, allowing the agent to navigate spaces with asymmetric transitions. When presented with single-unit input from the *goal* layer, activity within *goal proximity* filters through the recurrent connections, losing strength at each jump and ultimately forming a gradient of activity representing the proximity of each state to the goal. This process is roughly analogous to a full backward sweep from the goal, except that all paths are explored simultaneously. At the same time, the activity in *adjacent states* constitutes a one-state simultaneous forward sweep from the current state. The model essentially uses gradient ascent to move from the current state closer and closer to the final goal, similar in broad strokes to other approaches utilizing gradient ascent to maximize reward or achieve a goal (e.g., Friedrich et al., 2011; Legenstein et al., 2010).

### Neural Models

To our knowledge, the GOLSA model is the first to illustrate how flexible goal-directed learning can be achieved in a biologically plausible continuous-time neural network, with plasticity driven solely by local information during behavior as opposed to a dedicated learning period. However it is instructive to compare it to alternative neural network models dealing with similar issues. Of particular interest is how these networks deal with the dual roles problem, in which the same connection that drives actions based on desired consequences must learn based on observed action-consequence pairs.

Baldassarre et al. (2013) describe a model containing three basal ganglia (BG)-cortical loops governing arm actions, visual attention, and goal selection. Each loop has a BG/thalamic circuit which feeds into a cortical component that projects to other parts of the network. While outputs from the arm and eye loops project outside of the model to the agent “body,” the goal loop output innervates the cortical components of the other two loops via a learned all-to-all connection. This allows the goal loop to override the novelty-driven “bottom-up” actions that the action loops would select in the absence of goal loop input.

Early in the learning phase, saccade location and arm actions are taken at random. When an unexpected outcome is witnessed, such as a box opening, a burst of dopamine (DA) triggers several simultaneous learning processes. During the learning phase, environmental outcomes activate the corresponding cortical units in the goal loop. The presence of DA strengthens the cortico-cortical connections between the goal loop output and the units representing the recently taken actions in the arm and eye loop cortical second layers. In the future, when an outcome unit is activated it will tend to activate the associated actions. At the same time, novelty drives repetition of efficacious actions until the results are no longer surprising, facilitating efficient exploration of the state transition structure. During the test phase, units in the goal loop are activated based on desired outcomes, rather than observed outcomes, which in turn activate the appropriate associated actions.

Despite the somewhat different emphases of the GOLSA model and the model described above, they each present a neural architecture for goal-directed learning and action. One relative strength of the Baldassarre et. al model is the ability to draw a tight correlation between some model components and descriptions of actual circuits at a systems neuroscience level. However, the architecture produced by this method exhibits several shortcomings relative to the GOLSA model as an account of goal-directed behavior generally. Due to the nature of the task it was constructed for, the model is capable only of one-step transitions toward a goal. Opening a box requires simply pressing the appropriate button while looking at it, modeled as two simultaneous actions, and there is therefore no sense in which the model can move closer to a goal without achieving it.

Since this model, like the GOLSA model, involves learning action consequences and then later activating those consequences as goals in order to drive the appropriate actions, it must also deal with the dual roles problem. Though this problem is rarely discussed as such, every mechanistic account of associative goal learning must solve it in some fashion, often in an implausible way. In this case, the goal loop performs two fundamentally different roles in fully segregated learning and test phases. The biologically implausible nature of this solution is somewhat masked by the presence of explicit learning and test phases in the real-world button-box task, whether the task is performed by the model or animals. The architecture would require substantial revision to handle cases which lack this structure, including naturalistic exploratory and goal-directed behavior.

Given the extraordinary utility of reinforcement learning for problems similar to those faced by animals, RL offers a promising theory of the computational process underlying goal directed behavior. However, it is challenging to demonstrate how RL can be carried out in a neural substrate. Whereas the GOLSA model and the model of Baldassarre et al. (2013) approach goal directed learning with a somewhat different strategy than pure RL, Friedrich and Lengyel (2016) purport to show how RL methods can be embedded in spiking neurons. In their model, neurons represent state-action (SA) conjunctions. If a neuron represents an action from state j, the strength of the projection to all other neurons is equal to the probability that the target unit’s SA pair will lead to state j. Units representing alternative actions in the same state are mutually inhibitory. Activity is seeded into the network via a reward unit projecting to all units with strength equal to the amount of reward that can be attained from the represented SA pair.

The reward function R(s, a) is represented by the weights between the reward unit and the SA units. The transition function T(s, a, s’) is represented by the weights between units representing each (s, a) pair and all units representing (s’, a’) pairs for every action a’. Finally, the values of SA pairs, Q(s, a) are represented as the steady state activity of each unit after the network dynamics settle. Because the model replicates MBRL, it is capable of using the transition information to flexibly switch between different goals by replacing the reward neuron with on whose weights specify which SA pairs lead to the new goal.

This spiking RL model combines the backward diffusion of activity from the goal found in the *goal-gradient* layer of the GOLSA model with the action selection components by diffusing activity through state action pairs rather than states alone. Thus, the model appears to offer a much-simplified architecture with the same capabilities as GOLSA. In particular, the model appears to completely sidestep the dual roles problem without any need for an oscillatory mechanism or distinct learning and acting phases.

However, the devil is in the details of the learning procedure. The excitatory transition weights are updated by a learning law which increases the strength of the weight on unit i from unit j if the agent completed state-action pair i and is now in state j. In the future, reward activity propagating from the reward unit to unit j will activate unit i, making it more likely for the agent to take action i in state i.

A major problem with this scheme is that the network has no plausible method of determining its past and current state, yet knowledge of the current state is required in action selection, since actions are selected by comparing the values of the neurons representing various actions that can be taken in the current state. Fundamentally, this problem stems from the fact that whereas mathematical variables can be defined at will, there are relatively few neural properties (and even fewer neural properties that are typically modeled) that can be used to encode values. Typically, as is the case here, those properties are firing rate and weight strength. Since the above model uses only a single population of neurons with weight strength encoding transition probability and firing rate encoding value, there is no representational space left over for the current state and previous state. Because of this, learning relies crucially on non-neural signals about critical information (rather than mere control signals), which are difficult to interpret biologically. Furthermore, synaptic changes are the result of presynaptic and postsynaptic firing but in this case firing and plasticity are totally divorced since synaptic strength is based on transition probability while firing is governed by value.

In a very similar fashion to the GOLSA model, Hasselmo et al. (2005) address the dual roles problem utilizing an oscillatory scheme, derived from neurophysiological work investigating how rat hippocampus and PFC cooperate to support navigation (Hasselmo et al., 2002). Despite the use of a similar oscillatory scheme, the models are dissimilar in their level of analysis and overall structure. In the GOLSA model, each layer performs a particular function while individual units represent particular states, actions, and state transitions. In Hasselmo’s model, each state and action is associated with a subnetwork, representing a cortical minicolumn.

Each minicolumn contains a miniature version of *goal-gradient* and *adjacent-states*, linking actions and states which are one transition away. The activity of all minicolumns collectively generate the total gradients. Goal activity propagates backwards from a state minicolumn representing the goal, while information about consequences propagates forwards from the current state. As presented, the model does not make full use of the forward propagation, though the authors state that future work could utilize the mechanism to plan forward trajectories.

## Limitations and Possible Extensions

Like any model, the GOLSA model is limited in a number of important ways. In its present form, the model can only navigate discrete state spaces. While this works for gridworlds and games with distinct states, many environments and situations in the real world cannot be reasonably approximated in a discrete fashion. The brain consists of discrete neurons that produce all-or-nothing action potentials, so it must have mechanisms in place to appropriately represent continuous spaces with a discrete substrate. How to do this at both the algorithmic and neural levels is an area of active research. One solution is to tessellate the continuous space into discrete regions, a solution employed in some varieties of RL (Barto, Sutton, & Anderson, 1983; Doya, 2000; Santamaria, Sutton, & Ram, 1996). Grid and place cells in the hippocampus seem to employ a variant of this strategy, though their representational power is significantly greater than binary vectors since a particular cell can respond in a graded fashion based on how close the agent is to the center of the cell’s spatial receptive field (Hafting, Fyhn, Molden, Moser, & Moser, 2005; O’Keefe & Dostrovsky, 1971).

Even if the state space can be discretized, in many cases there a large number of important states. This poses a problem for the one-hot encoding scheme used in the GOLSA model since most layers have either n or n^2^ units, where n is the number of states. There simply are not enough neurons in the brain to assign one to each important state, let alone every possible transition between states. The representational scheme used in the GOLSA model is quite inefficient since it limits vector representations to the standard basis vectors of the high dimensional space without utilizing any of the intermediate space. While this method was chosen for simplicity, other representational frameworks offer greater power. For instance the Neural Engineering Framework provides a method for embedding complex vector representations and operations in networks of spiking (or rate-coded) neurons (Eliasmith & Anderson, 2003; Stewart, Bekolay, & Eliasmith, 2011). Function approximation is a related approach in which a model function attempt to mimic a target function (like a reward or transition function), even if the properties or amount of units available are insufficient to match the target exactly. This method has been successfully used to adapt RL to continuous spaces (Doya, 2000; Millan, Posenato, & Dedieu, 2002; Santamaria et al., 1996).

Beyond the need for extremely large model layers, large state spaces pose a particular problem for the goal-gradient, which constitutes one of the central mechanisms in the GOLSA model’s operation. In fact, the goal-gradient is limited in several important ways. As modeled here, firing rates can only vary between 0 and 1, but for the goal gradient to function, all states must be represented in that space. As the unit activities get very close together, they become highly susceptible to noise interference. Similarly, the bias toward highly interconnected states would become more problematic. Recall that the activity gradient is established by a recurrent all-to-all connection within *goal-gradient* where each unit is connected to each other unit that represents an adjacent state. External activity is sent into one unit which then filters through the connections to the rest of the network. Units with more connections will receive more input and will therefore be more active than units with fewer connections that are the same distance from the goal. In a large state space, a unit representing a state 100 steps from the goal will be very nearly as active as a unit representing a state 99 steps from the goal. This small difference could be overcome by the interconnection bias.

Small differences between weight strengths could also substantially distort the gradient. In the simulations described in Chapter 2, the learning rates were set high enough that weights rapidly reached their upper limit and were thus equal across connections. For most projections this was not entirely necessary. The *adjacent-states* units, for example, only need to represent one of three facts about the state they represent, with three levels of activity: whether it is the current state (very highly active), whether it is an adjacent state (significantly active), or not adjacent (not active). Small differences in activity among the units representing states adjacent to the current state are not relevant for task function. The goal-gradient, in contrast, represents important information with small differences in activity. Furthermore, each weight in in the projection from *state* to *adjacent-states* is “used” once to weight the incoming activity. In the *goal-gradient*, each weight is “used” many times to weight recurrent activity. For example, unit 5 receives input from unit 4, which is affected by the weight between the two. Unit 5 sends input to unit 3, which in turn sends input to 4 which again gets filtered through the weight between units 4 and 5.

The requirement that the weights be uniform in the *goal-proximity-layer* is one of the reasons that only positive weight changes are allowed. This is obviously a limitation of the model, since it means that it cannot easily adapt to a changing environmental topology (though it can respond appropriately to changing drives or goals). “Unlearning” would pose several challenges to the framework presented here. First, when should a weight be eligible for a decrease in strength? In other words, what constitutes evidence that two states are no longer adjacent? A crude solution would be to have weights decay slowly over time, as if the agent continually forgot the structure of the environment. If experience were diverse and thorough enough, it could continually reset the decaying weights unless the environment really had changed. However, a slow decay would result in widely varying weight strengths which would distort the gradient.

A further limitation of all-or-nothing weights that code for adjacency is that they are difficult to adapt to situations in which state transitions are probabilistic rather than deterministic. Probabilistic state transitions are a major feature of Markov Decision Processes, the mathematical framework in which RL is formulated. This feature allows environments with some type of risk/reward tradeoff. For instance, in some state action A may yield a large reward rarely but otherwise nothing, whereas action B will yield a small reward consistently. An RL framework is able to weight the potential outcomes by their probabilities and thereby make an informed decision. In contrast, GOLSA as currently formulated is incapable weighting outcomes by their probabilities. It selects a desired goal state and then uses the knowledge of the state space to pursue it. On the one hand, this is a large advantage over RL, since it can pursue an arbitrary goal state rather than a unitary reward signal. However, in the situation described above, the GOLSA model would rather mindlessly attempt to attain the maximum reward if that were selected as the final goal state, despite the risk.

The GOLSA model could be adapted in a number of ways to address these shortcomings. A more sophisticated goal-selection procedure than the simple drive-based system would be needed for situations in which the agent may not be able to reach a desired goal state on a particular trial due to the stochastic nature of the task. The additional mechanisms would need to be able to weight the magnitude of each outcome’s potential drive reduction (i.e., the value) by the probability of attaining that outcome. In fact, this could be a place where a system very similar to RL could be integrated into the GOLSA model, since RL methods have already proven successful at dealing with these types of tasks.

The challenge posed by large state spaces may be overcome by nesting models. The weight matrices for the projection from *state* to *adjacent-states* and within *goal-gradient* essentially constitute maps of the topology of the environment. In the case of real maps, no single paper map of the environment contains all the necessary detail for the whole world. Distinct maps capture the layout of the world at differing levels of abstraction and resolution. A high-abstraction map is useful for making a general plan, while a low-abstraction, high resolution map is useful for working out the details within a particular region of the larger map. There is not enough space on a reasonably sized paper map to capture all of the detail over a large area. Likewise, there is not a wide enough range in activation levels to adequately represent a very large state space. Therefore, one promising future direction for the GOLSA model would be the introduction of nested maps of the environment. The high-level portion of the model would select which sub-region of the entire space that the agent should move toward, which would in turn specify a low-level goal like a door leading from one sub-region to another. Nested models have been successfully applied in Q-learning (Digney, 1996), and parts of prefrontal cortex appear to exhibit a nesting structure as well (Koechlin, Ody, & Kouneiher, 2003).

## Acknowledgments

We thank A. Ramamoorthy for helpful discussions and J. Fine for helpful feedback on the manuscript. JWB was supported by NIH R21 DA040773.

