## Supplementary material for "Computational neural mechanisms of goal-directed planning and problem solving"

**Supplementary Materials**

### Model Components

#### Layers

The GOLSA model is constructed from a small set of basic components. The main component class is a layer of units, where each unit represents a neuron (or, more abstractly, a small subpopulation of neurons) corresponding to either a state, a state transition, or an action. The activity of units in a layer represents the neural firing rate and is instantiated as a vector updated according to a first order differential equation (c.f. Grossberg, 1973). The activation function varies between layers, but all units in a particular layer are governed by the same equation. The most typical activation function for a single unit is,

$$da(t) = \left(-\lambda a(t)dt + \left(1 - a(t)\right)Edt - Idt + \varepsilon N(t)\sqrt{dt}\right), \qquad \text{(S1)}$$

where *a* represents activation, i.e. the firing rate, of a model neuron. The five terms of this equation represent passive decay, shunting excitation, linear inhibition, and random noise. "Shunting" refers to the fact that excitation (E) scales inversely as current activity increases, with a natural upper bound of 1. The passive decay works in a similar fashion, providing a natural lower bound activity of 0. The inhibition term linearly suppresses unit activity, while the final term adds normally distributed noise ($\mu$=0, $\sigma$=1) . Because the differential equations are approximated using the Euler method, the noise term is multiplied by $\sqrt{dt}$ to standardize the magnitude across different choices of dt (Busemeyer & Townsend, 1993; Usher & McClelland, 2001). The speed of activity change is determined by a time constant $\tau$. The parameters $\tau$, $\lambda$, $\varepsilon$ vary by layer in order to implement different processes. E and I are the total excitation and inhibition impinging on a particular unit for every presynaptic unit j in every projection p onto the target unit,

$$E = \sum_p \sum_j [w_{p_j} a_{p_j}]^+ \tag{S2}$$

$$I = \sum_p \sum_j [w_{p_j} a_{p_j}]^- \tag{S3}$$

A second activation function used in several places throughout the model is,

$$da(t) = \left(-\lambda a(t)dt + \left(1 - a(t)\right)Edt - a(t)Idt + \varepsilon N(t)\sqrt{dt}\right) \tag{S4}$$

This function is identical to Equation (6) except that the inhibition is also shunting, such that it exhibits a strong effect on highly active units and a smaller effect as unit activity approaches 0. While more typical in other models, shunting inhibition has a number of drawbacks in the current model. Two common uses for inhibition in the GOLSA model are winner-take-all dynamics and regulatory inhibition which resets layer activity. Shunting inhibition impedes both of these processes because inhibition fails to fully suppress the appropriate units, since it becomes less effective as unit activity decreases.

**Projections**

Layers connect to each other via projections, representing the synapses connecting one neural population to another. The primary component of projections is a weight matrix specifying the strength of connections between each pair of units. Learning is instantiated by updating the weights according to a learning function. These functions vary between the projections responsible for the model learning and are fully described in the section below dealing with each learning type. Some projections also maintain a matrix of traces updated by a projection-specific function of presynaptic or postsynaptic activity. The traces serve as a kind of short-term memory for which pre or postsynaptic units were recently activated, which serve a very similar role to eligibility traces introduced in Barto et al. (1979), though with a different mathematical form.

**Nodes**

Nodes are model components that are not represented neurally via an activation function. They represent important control and timing signals to the model and are either set externally or update autonomously according to a function of time. For instance, sinusoidal oscillations are used to gate activity between various layers. While, in principle, rate-coded model neurons could implement a sinusoidal wave, the function is simply hard coded into the update function of the node for simplicity. In some cases, it is necessary for an entire layer to be strongly inhibited when particular conditions hold true, such as when an oscillatory node is in a particular phase. Layers therefore also have a list of inhibitor nodes that prevent unit activity within the layer when the node value meets certain conditions. In a similar fashion, some projections are gated by nodes such that they allow activity to pass through and/or allow the weights to be updated only when the relevant node activity satisfies a particular condition. Another important node provides strong inhibition to many layers when the agent changes states.

**Environment**

The agent operates in an environment consisting of discrete states, with a set of allowable state transitions. Allowable state transitions are not necessarily bidirectional, but they are deterministic (unlike the typical MDP formulation used in RL). In some simulations, the environment also contains different types of reward located in various states, which can be used to drive goal selection. In other simulations, the goal is specified externally via a node value.

**Complete Network**

Each component and subnetwork of the model is described in detail below or in the main text, but for reference and completeness a full diagram of the core network is shown in Figure S1. Some of the basic layer properties are summarized in Table S1. Layers and nodes are referred to using italics, such that the layer representing the current state is referred to simply as *current-state*.

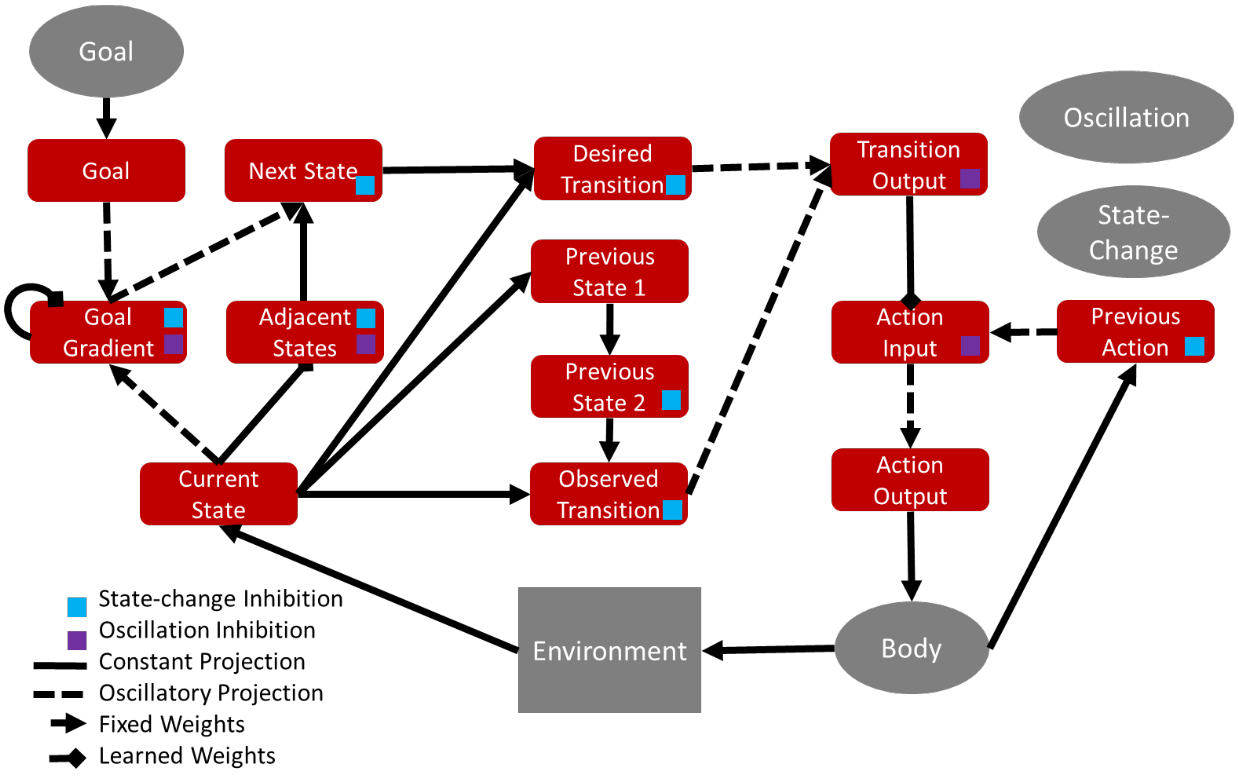


*Figure S1.* Full diagram of core model. Each rectangle represents a layer and each arrow a projection. The body is a node, and two additional nodes are not shown which provide inhibition at each state-change and oscillatory control. The colored squares indicate which layers receive inhibition from these nodes. Some recurrent connections not shown.

| Layer Name | Inhibition Type | Time Constant | Decay Rate | Noise Gain |
|---|---|---|---|---|
| current-state | shunting | 0.5 | 1 | 0 |
| goal | linear | 1 | 1 | 0 |
| goal-gradient | linear | 1 | 1 | 0 |
| adjacent-states | linear | 0.2 | 1 | 0.01 |
| next-desired-state | linear | 1 | 0.1 | 0 |

| | | | | |
|---|---|---|---|---|
| previous-state-1 | shunting | 4 | 0.5 | 0 |
| previous-state-2 | linear | 1 | 0.001 | 0 |
| previous-action | linear | 1 | 0.001 | 0 |
| observed-transition | linear | 1.5 | 1 | 0 |
| desired-transition | linear | 1.5 | 1 | 0 |
| transition-output | linear | 1.5 | 1 | 0 |
| action-input | linear | 1 | 1 | 0 |
| action-output | linear | 0.5 | 0.2 | 0 |

*Table S1. Full list of model layers and associated parameters.*

### Model Operation

#### Determining Desired Next State

The first two steps in the model's goal-pursuit "algorithm" determine which state to try to navigate to next (Figure S2). This process utilizes a series of layers each containing n units where n is the number of states in the state space, representing each state with a localist code (i.e., "one hot", or one unit per state). These layers work together to determine which states are currently possible to transition to and which of these states will take the agent closer to the goal. The subnetwork that accomplishes this process is illustrated in Figure S2.

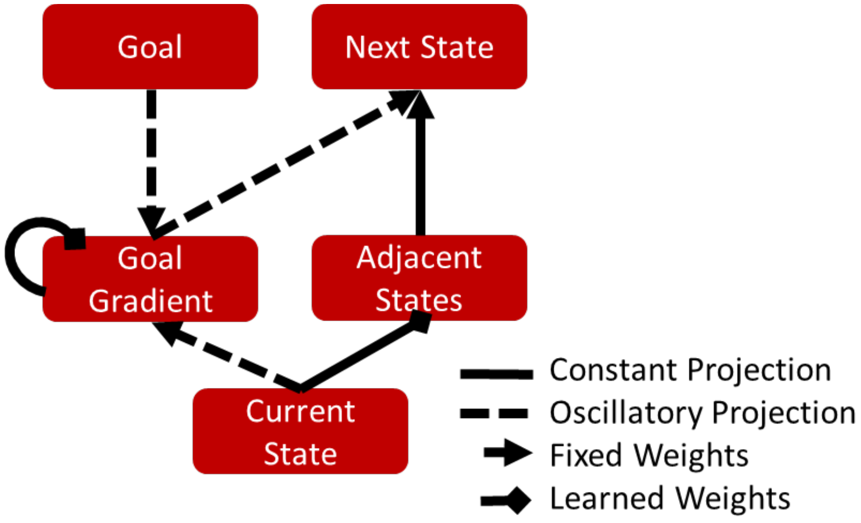


*Figure S2.* Subnetwork of layers responsible for determining the next desired state.

The *current-state* layer receives input directly from a node representing the environment, specifying a single state that the agent is in. *Current-state* is connected to the *adjacent-states* layer via an all-to-all learned projection. After learning, *adjacent-states* exhibits strong activity in each unit representing a state that could be reached from the current state via one transition, including the current state itself. The states represented this way form the candidates for the desired next state.

A desired next state is chosen from these candidate states using activity from the *goal-gradient* layer. This layer is recurrently connected to itself via an all-to-all learned projection. For every possible state transition between a starting state s and a destination state s', (s, s'), the final learned projection exhibits a connection between the unit representing s' and the unit representing s. This allows externally injected activity to "flow backwards" through time, activating states further and further from the state represented by the unit receiving the initial injection of input.

The key input to this layer comes from a layer representing the desired goal, again in a localist fashion with a single unit representing each state. When this activity is fed into the *goal-gradient* layer, recurrent connections between units within the layer facilitate the spreading of activation from the unit representing the goal to units representing nearby states. Because excitation scales inversely with the current level of activation, recurrent activity diminishes as it passes through each model synapse, establishing a gradient of activity with maximum activity at the unit representing the goal and diffusing outward. The activity in this layer spreads to units representing states that precede the goal state.

The *next-desired-state* layer uses lateral inhibition and recurrent excitation to create a WTA competition. When inputs to this layer differ even slightly in activation, the differences are quickly amplified such that a winning unit achieves a high level of activation while all other units exhibit almost no activation. The key input to this layer in is the *goal-gradient* layer which connects one-to-one. If all units from *goal-gradient* provided input to *next-desired-state*, the unit representing the goal state would always win the competition in *next-desired-state* since the peak of the gradient is always the goal state. Instead, the network needs to use the gradient to discriminate among only the states adjacent to the agent's current location. To achieve this, the projection from *goal-gradient* to n*ext-desired-state* is gated on a unitwise basis by activity from *adjacent-states*. In other words, *adjacent-states* specifies which units in are allowed to compete to be the next desired state, while *goal-gradient* distinguishes between them. The unit that will win the competition in *next-desired-state* will therefore be the unit representing the adjacent state that takes the agent closest to the goal. If there are multiple such states, the tie can usually be broken by the network's bias for more interconnected states (as detailed in the section on learning), though random noise can also be used.

Figure S3 below demonstrates the activities of the *current state, adjacent-states, goal-gradient*, and *next-desired-state* layers over the course of a single run starting at state 1 with the goal at state 6.

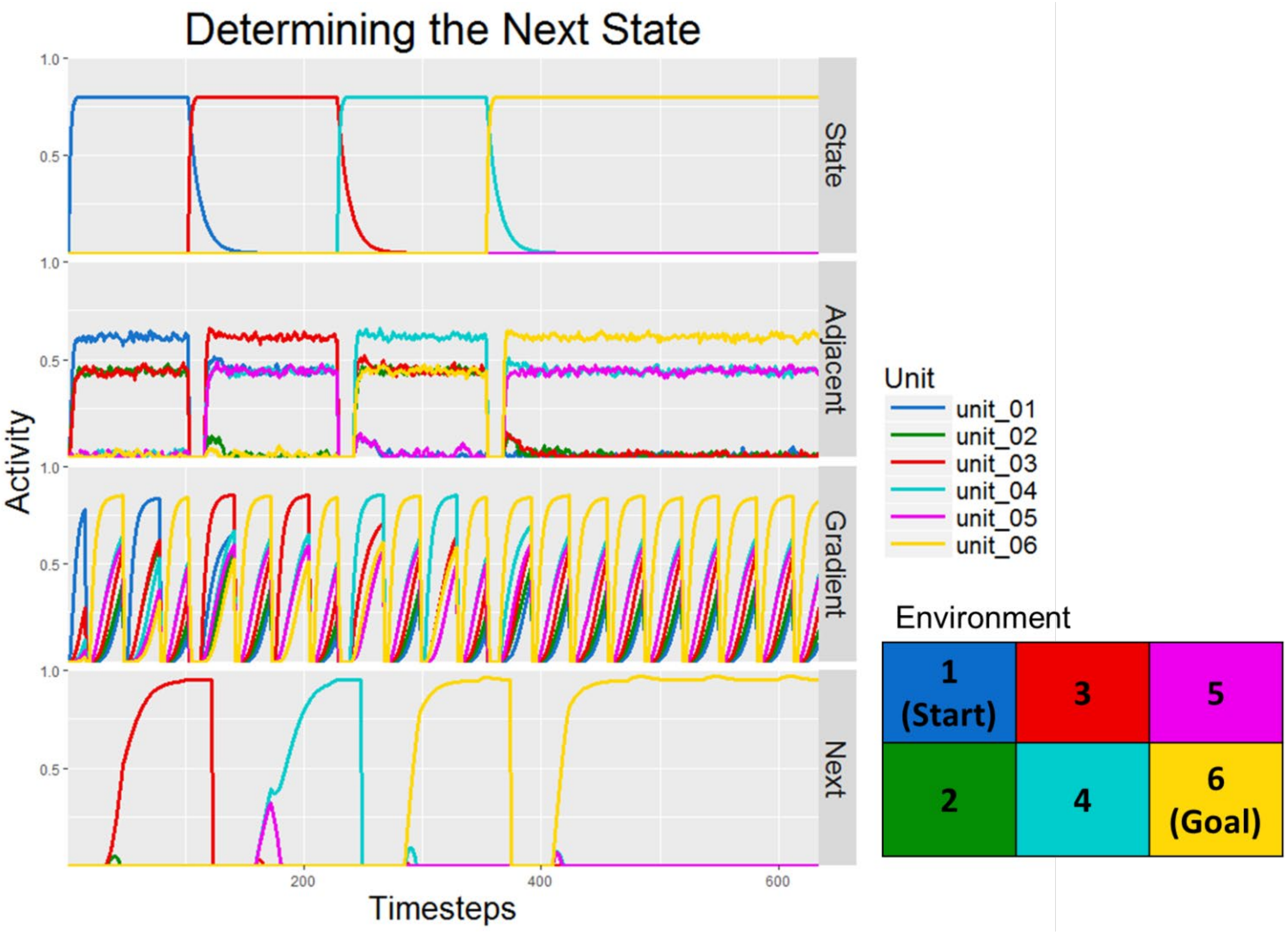


*Figure S3.* Activity plots for the current-state, adjacent-states, goal-gradient, and next-desired- state. Each unit represents the corresponding state. The State layer represents the current location of the agent in the environment shown. In the Adjacent layer, the unit corresponding to the current state is most active, but all units representing adjacent states are active as well. The Gradient layer oscillates between input from the goal layer (not shown) and the state layer. For each input, units activate in proportion to the distance between the state they represent and the input state. The Next layer uses input from Adjacent and Gradient to select the adjacent state which will bring the agent closer to the goal.

The *state* activity reveals three state transitions, first from state 1 to state 3, then 4, and finally the 6, the goal location. At each step, *adjacent-states* exhibits strongest activity for the state the agent is currently in, less activity for every state the agent is adjacent to, and 0 activity for the remaining states. The *goal-gradient* activity (Figure S4) is more complex. As explained in more detail below, learning requires that this layer alternate between receiving input from the current state layer during the learning phase and input from the goal layer during the acting phase of an oscillation. The gradient only informs the WTA competition in *next-desired-state* during the acting phase when the gradient peak is at the goal. The gradient can be seen more clearly in the Figure S4 below, showing the activity during a single acting phase of the oscillation.

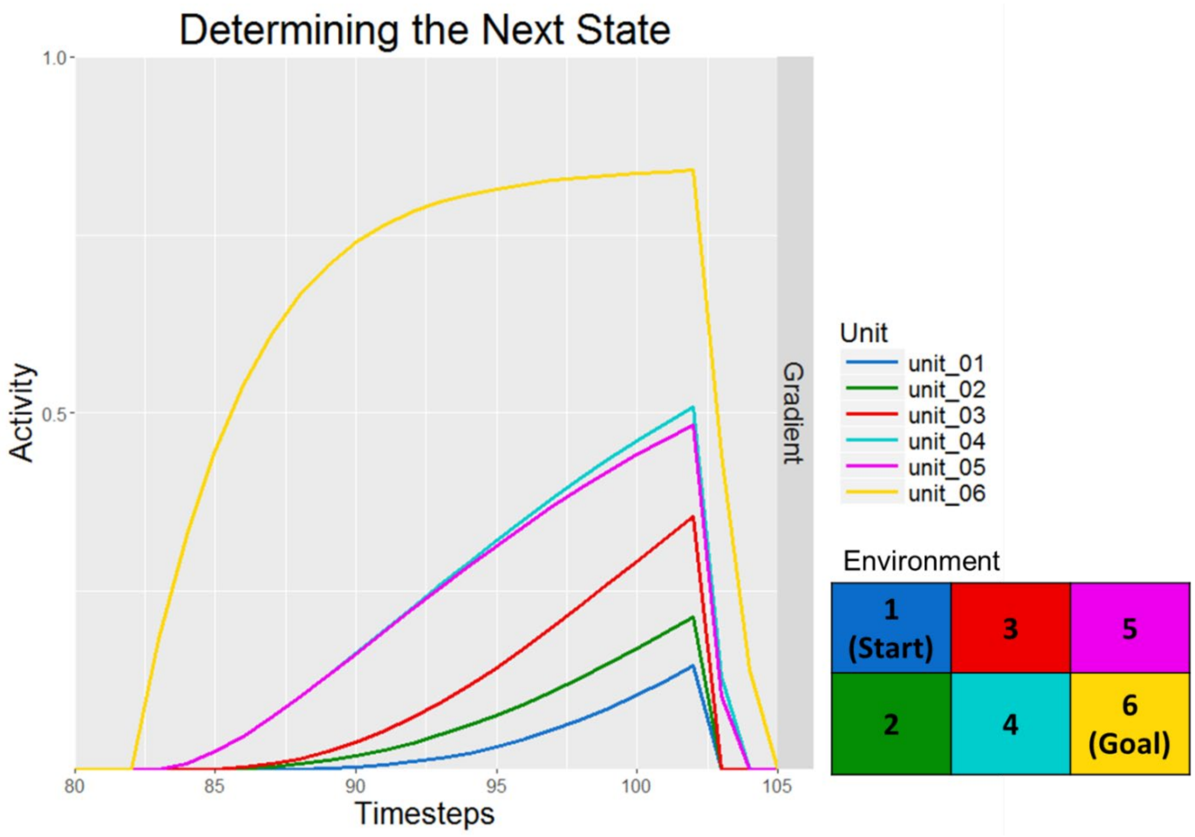


*Figure S4.* Goal gradient activity during one oscillatory cycle. The strongest activity is in unit 6, representing the goal state. Units 4 and 5 have the next strongest activity (4 is slightly more active because it is connected to more states – see the learning section for details). The current state of the agent is irrelevant for the activity of this layer when receiving input from the goal layer as in this figure.

As expected, each unit is active in rough proportion to how close it is to the goal state. Units 3 and 4 have an additional advantage over the others, due to their greater connectivity within the layer. After learning each unit is connected to every unit representing an adjacent state, so 3 and 4 receive extra excitation in virtue of their higher number of adjacent states. This feature actually helps break symmetry in the case of ties (for example in the first step), but biases the GOLSA agent toward states that are highly interconnected.

As shown in the bottom panel of Figure S3 , the *next-desired-state* layer implements a winner take all competition among inputs from *goal-gradient* gated by the activity in *adjacent-states*. The small bumps in the activity of multiple units shortly following the cessation of the inhibition reflect the brief period of competition before the layer activity settles. In some cases this process can take a bit longer as in the second step where the model was uncertain about whether to move to state 4 or 5 next, ultimately settling on state 4 due to the bias explained above.

Like most of the layers throughout the model, all layers other than *current-state* in Figure S3 are subject to a burst of inhibition following a state transition. This allows activity patterns to reset appropriately, especially layers with WTA dynamics as these layers are particularly resistant to change.

**Determining the Next Action**

The desired next state in combination with the observed current state specifies a desired state transition, represented by the *desired-transition* layer containing $n^2$ units where n is the number of states in an environment. The first n units represent transitions starting from the first state, the next n units represent transition starting from the second state, etc. Of course, not all of these units will represent possible transitions in a given environment. Projections from *current* *state* and *next-desired-state* contain hard-coded weight matrices which uniquely activate a *transition* unit for each combination of current and desired states. The *desired-transition* layer projects to the *transition-output* layer which in turn projects to an *action-input* layer containing one unit for each possible action. The basic architecture of this section of the network is depicted in Figure S5 below. The two transition and action layers facilitate oscillatory learning, described in more detail in the following section.

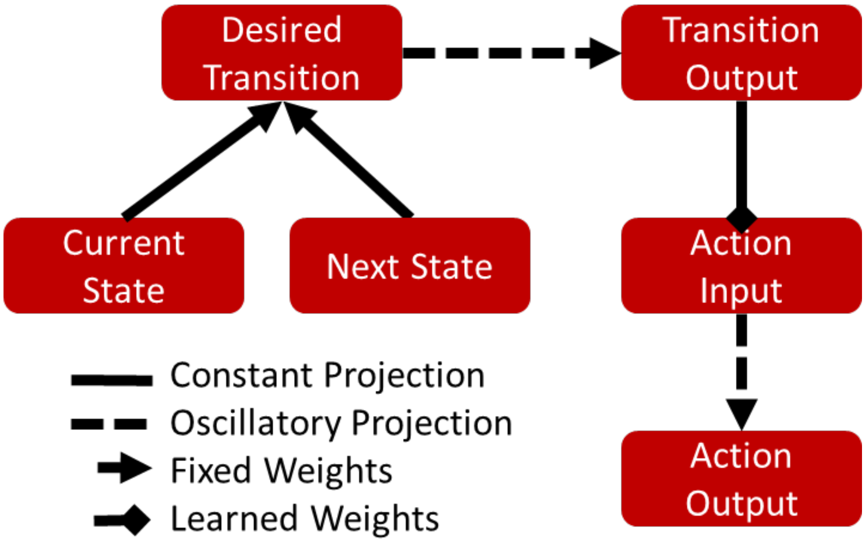


*Figure S5.* Subnetwork for selecting an action. The current and next states specify a desired transition which feeds into the action system. The importance of multiple transition and action layers is explained in the section on learning below.

After learning, each transition is mapped to the appropriate action, implementing the desired state transition as specified by the current and next desired states. The cycle then begins again from the new state location until the agent has reached the goal. Figure S6 below shows the activity of the *current-state*, *next-state*, *desired-transition* (abbreviated 'TransDes'), and *action-out* layers over the course of the same simulation as previous figures.

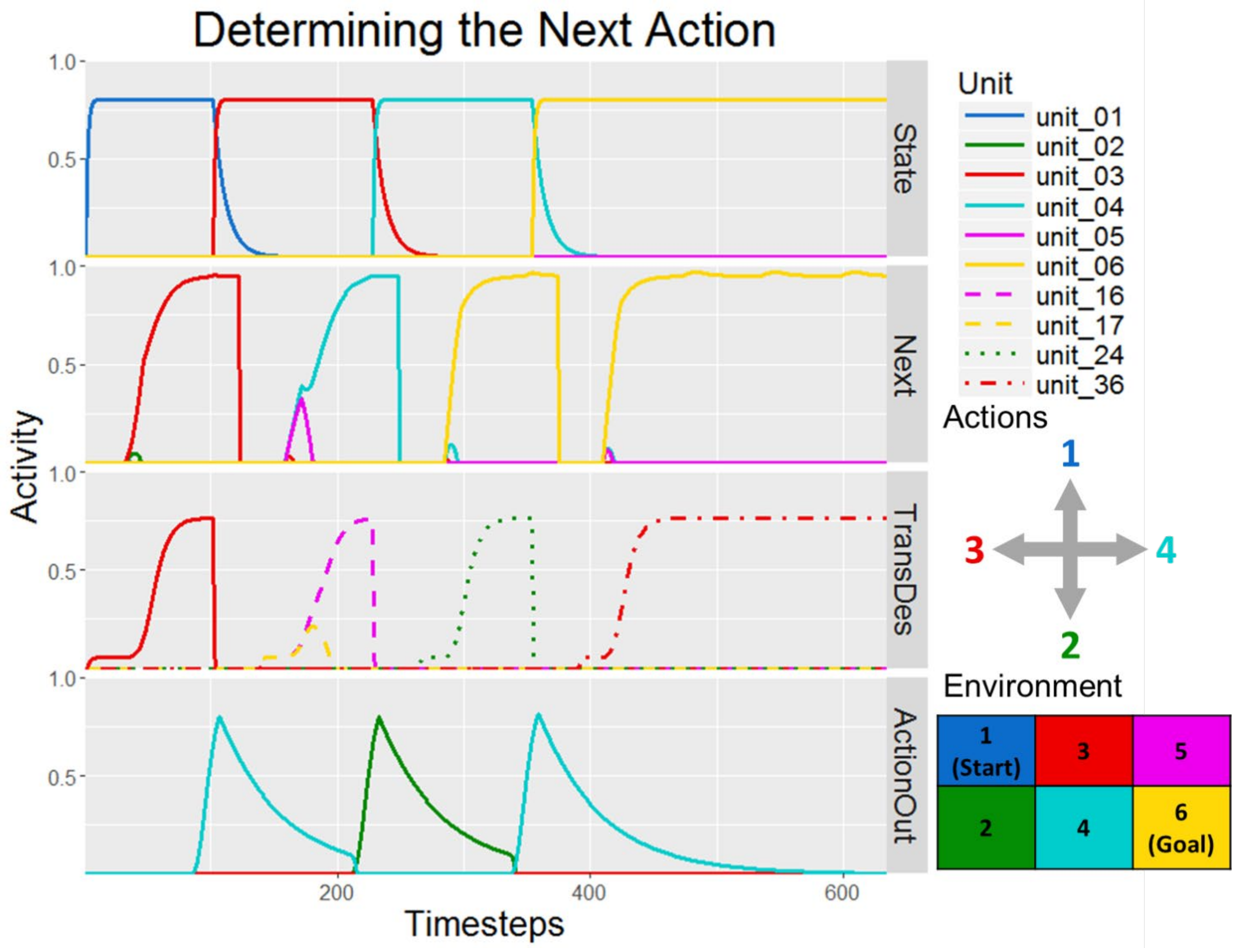


*Figure S6.* Activity plots for several layers during a trial starting in state 1 with the goal in state 6. Note that of the layers displayed here, only State and Next code for states. Desired Transition encodes transitions (of which there are 36) and Action Out encodes actions (of which there are 4).

Each *current-state* and *next-desired-state* pair activates the correct transition which in turn activates the appropriate action (move right, move down, move right). Activity in *action-out* above a threshold activates a node representing the agent's body (not shown) which implements the action and enforces a delay before the next action can be taken.

The *desired-transition* and *action-out* layers sit on the outside of the main action-learning circuit and are both concerned solely with action. Intermediate between these layers are the *transition-output* and *action-input* layers which alternate between an acting and learning phase, similar to the *goal-gradient* layer, as shown in the following section.

To this point, we have seen how the GOLSA functions after learning. It is comprised of a series of layers of model neurons, where each layer dedicates one neuron to representing a state, a transition, or an action. Four layers represent the current state, adjacent states, the goal state, proximity of the states to a chosen goal. An additional layer represents the next desired state which is determined by selecting the adjacent state which brings the agent closer to the goal. Once the next desired state is selected, the network drives the action which brings about the desired transition. In order to do this, the network must accurately represent the (forward-in-time) topology of the network in the weights between *state* and *adjacent*, as well as the (backward-in-time) topology in the weights from *goal-gradient* to itself. In addition, it must specify the correct mapping from desired transitions to actions in *transition-out* to *action-input*. All three can be learned via exploration using biologically plausible learning laws.

### Learning in the Core Model

Learning in the core GOLSA model takes place in three projections: from *current-states* to *adjacent-states*, from *goal-proximity to itself,* and from *transition-output* to *action-input*. Each rely on modified Hebbian learning and require relatively little experience in the environment as described below.

**Current-state to Adjacent-states**

Units in the *adjacent-states* layer should become highly activated when receiving input from a *current-state* unit representing a state that directly precedes the state represented by the *adjacent-states* unit (see Figure 12). Therefore, when the agent moves from state 1 to state 2, the weight between the presynaptic state 1 unit and the postsynaptic state 2 unit should increase.

After each transition, there will be high postsynaptic activity in the *adjacent*-states unit corresponding to state 2 since the connection from *current-state* to *adjacent-states* is initialized at one-to-one, but the presynaptic state 1 unit will no longer be active. Eligibility traces are therefore necessary in the presynaptic component of the learning law. All eligibility traces in the core model are of the form,

$$\dot{r} = \begin{cases} (r-1), & a_{pre} < \rho \\ 1, & a_{pre} \geq \rho \end{cases} \quad \text{(S5)}$$

When presynaptic activity, $a_{pre}$, meets the threshold ρ (in this projection ρ = .6), the trace increases to a maximum of 1 while if presynaptic activity is less than ρ, the trace decreases slowly at first and then more quickly, to a minimum of 0. After each transition, the trace from the previously activated state will still be high at the same time that the newly reached state unit is highly active.

Once the weights are learned, incoming activity from state 1 will activate all units representing states that could be reached in one transition from state 1. This would cause an issue with the learning scheme described above since now learning would occur between the previously active *current-state* unit and all of the now active states adjacent to state 1. To solve this problem, the learned weights are capped at 1, while the one-to-one initial weights have a strength of 2. As a result, the adjacent states layer exhibits a pattern of activity in which the unit corresponding to the current state is most active while all adjacent units are active to the same, but lesser degree (see Figure 3). This pattern can be thought of as a highly truncated proximity map from the current location, with only three weight values of import, namely 0, 1, and 2. The discrete nature of this scheme is reflected in the learning law,

$$\dot{w} = \eta\left[\left(\left(r_{pre} > \chi\right) * \left(a_{post} > \varphi\right)\right) - w\right]^{+} \tag{S6}$$

where the inequality evaluates to 1 if true and 0 otherwise, and the values $\chi = .7$ and $\phi = .6$ were chosen as reasonable thresholds. The role of $\chi$ is to isolate the trace representing the previously active state while the role of $\varphi$ is to isolate the current-state signal in *adjacent-states* activity which is more active than units representing the adjacent states.

Initially, the weights between corresponding *current-state* and *adjacent-states* are initialized to a strength of 2. After a single step in the first trial, from state 1 to state 2, the weights are updated to reflect the new knowledge about the environment, as unit 1 in *state* becomes strongly connected to unit 2 in *adjacent-states.* This process continues through the trial, and the final weight configuration after many trials accurately reflects the topology of the state space. This process is illustrated in Figure S7 below.

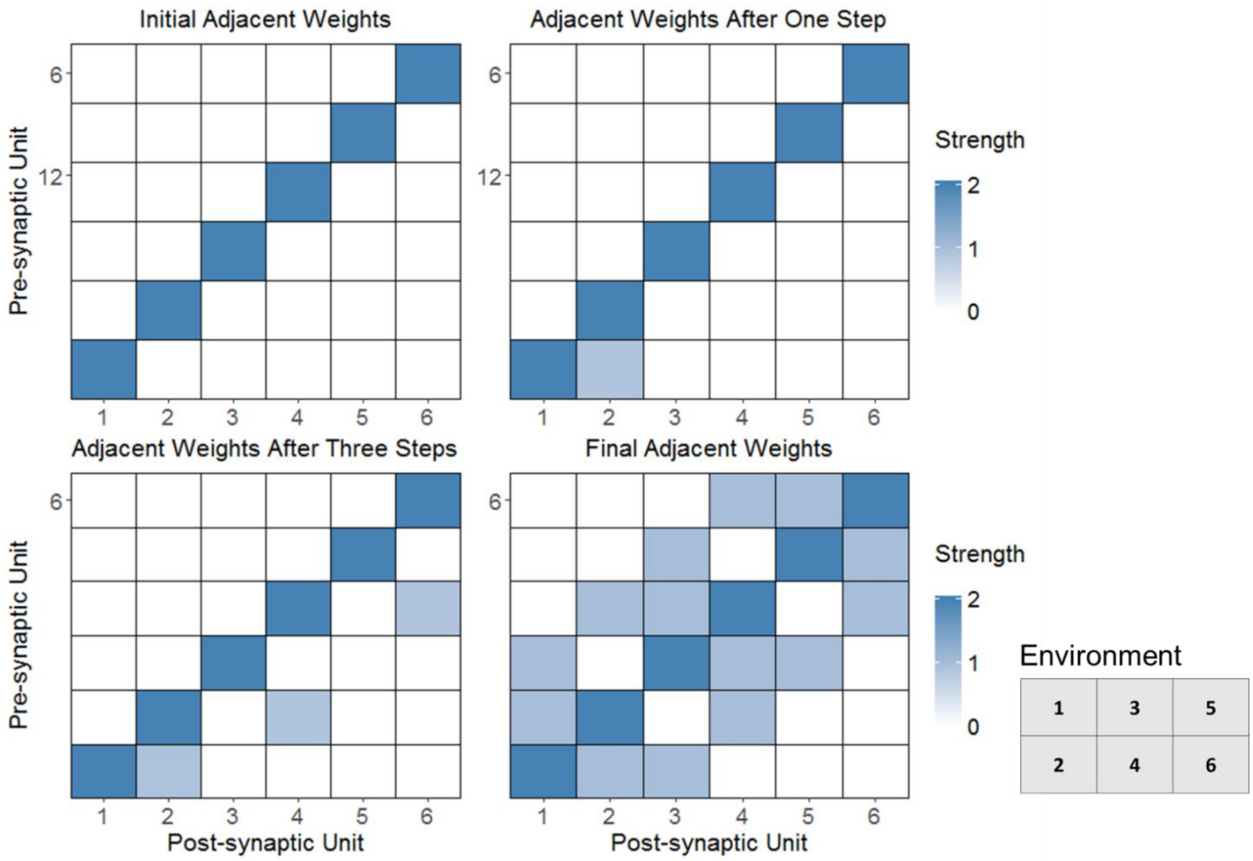


*Figure S7.* Progression of weights between state and adjacent-states starting with the initial values in the upper left, the weights after one step (from state 1 to state 2) in the upper right, the weights after a full trial in the lower left, and the final configuration of weights after all learning in the lower right. For example, states 4, 5, and 6 can be reached from state 6 so unit 6 in state projects to units 4, 5, and 6 in adjacent-states. Note that weights are strongest along the diagonal which ensures that the adjacent-states unit corresponding to the current state is most active. This is an important signal for learning.

#### Goal-proximity to Goal-proximity

The weight matrix used to form the goal proximity gradient is closely analogous to that learned between *current-state* and *adjacent-states*. One major difference is that whereas the latter projects from the current state into the future, the goal proximity gradient works backward through time. After experiencing a transition from state 1 to state 2, therefore, the weight from the state 2 unit should increase its connection to the unit representing state 1 rather than vice versa. This reversal simply requires that the learning law take account of postsynaptic activity traces and presynaptic activity, rather than the reverse as in the projection from *current-state* to *adjacent-states*. Whereas the learning related to the adjacent states occurred in the projection from one layer to another, for *goal-gradient*, the learning takes place in the weights of a recurrent projection from *goal-gradient* to itself.

The learning objective is for each unit to be connected to units representing adjacent states with a strength of 1. If that target is achieved, then external activity targeting a single *goal-gradient* unit will spread throughout the layer, first to units representing adjacent states, then to units representing states two transitions away, etc. Due to the shunting excitation in the activation equation, the strength of activation diminishes at each step, thereby establishing the gradient.

The learning within *goal-gradient* also differs from that in *adjacent-states* in that the network learns based on input from *current-state* but only informs goal-pursuit based on input from *goal*. Input to *goal-gradient* must pull double duty by representing both the current state, for learning, and the goal state, for acting. The weights in the recurrent connection from *goal-gradient* to itself need to be updated based on different input than the input the weights need to transform during goal-directed action. This pattern occurs in several places throughout the network and is hereafter referred to as the *dual roles problem*.

Separate learning and acting network configurations (c.f. Baldassarre 2013) could solve the dual roles problem if *goal-gradient* received its input from *current-state* during the learning trials and from *goal* during the acting trials. However, cognitive agents are obviously capable of learning while pursuing goals. To allow simultaneous action and learning, inputs oscillate based on the phase of a node, *substitution-oscillation* which oscillates according to $v = \cos(2t)$, as in

Figure S8, where *t* represents time[1]. In effect, the acting and learning phases are compressed in time to allow both to proceed together while the agent acts in the environment. During the learning phase, input from *current-state* is substituted for input from *goal*, and the learning law is active. To prevent spurious output leading the agent to pursue the current state, the connection from *goal-gradient* to *next-desired-state* is blocked. During the acting phase, input from *current-state* is blocked, input from *goal* is allowed, learning is inactive, and the connection from *goal-gradient* to *next-desired-state* is opened. The gradient is relatively sensitive to erroneous activity which presents a problem for this oscillatory scheme since all activity from one phase must fully decay before the gradient can be properly established in the next phase. To deal with this, the *goal-gradient* layer receives strong inhibition in between the learning and acting phases, in addition to the inhibition experienced by almost every layer at each state transition. Oscillations in the theta band as well as other frequencies may play a similar role in the brain, serving as a clock for learning and inhibition (Buzsáki, 2009; Hyman et al., 2003)

[1] The simulations shown here use the Euler method to approximate the evolution of the differential equations governing the system, with dt = .05 seconds. The "timesteps" shown on the x axis of the activity plots above refer to each dt step and each unit of time (seconds) therefore includes 20 timesteps.

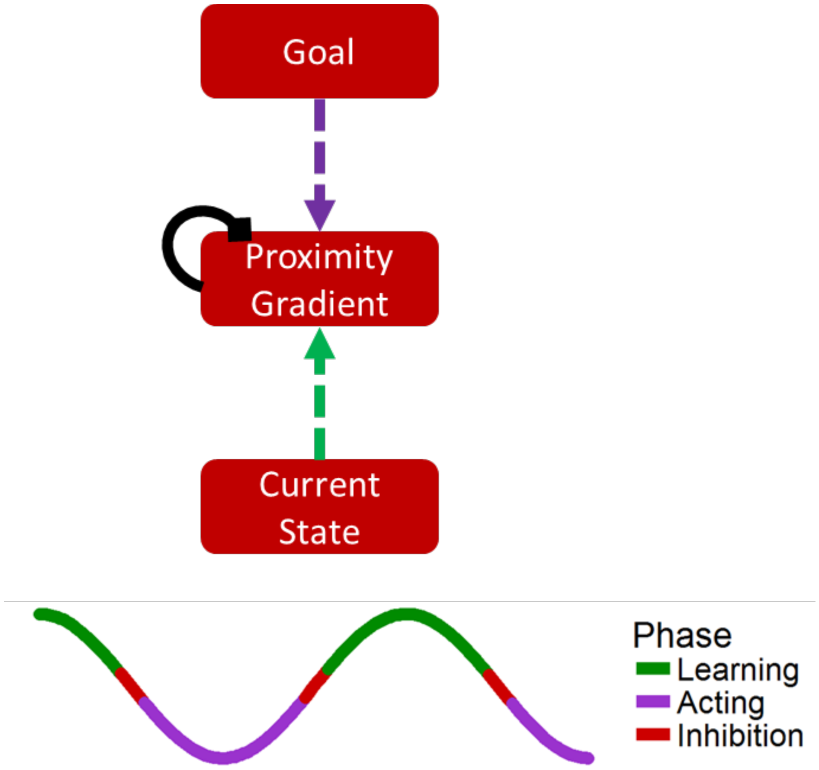


*Figure S8.* Network components involved in learning the goal-gradient weights. An oscillation controls alternating input from representations of the current state (for learning) and the goal state (for acting). Inhibition in between phases clears the activity in goal-gradient, ensuring that the gradient in each phase is not contaminated with remnant activity from the previous phase.

As with *adjacent-states*, unit activity above a fixed threshold (ρ= .6) in the learning phase indicates that the agent is in that state, and also starts the eligibility trace. After a state transition, the weights on the unit representing the previous state are eligible to learn, while presynaptic activity over a fixed threshold accounts for the other half of the learning law,

$$\dot{w} = \theta\eta\left[\left(\left(a_{pre} > \varphi\right) * \left(r_{post} > \chi\right)\right) - w\right]^{+} \qquad \text{(S7)}$$

where θ is 1 if *substation-oscillation* is in the learning phase, and 0 otherwise, and the other parameter definitions are the same as for Equation (11), the learning law for the projection from *state* to *adjacent-states*. For this projection, $\chi = .7$ and $\varphi = .75$. As before, the role of $\varphi$ is to select only the most active unit which, during the learning phase, will correspond to the current state. The purpose of $\chi$ is to allow learning only when the trace is at the level expected following one state transition (though, because the trace decay accelerates over time, this parameter is highly flexible since the trace will rarely be active at all after two state transitions).

The progression of weights is illustrated below,

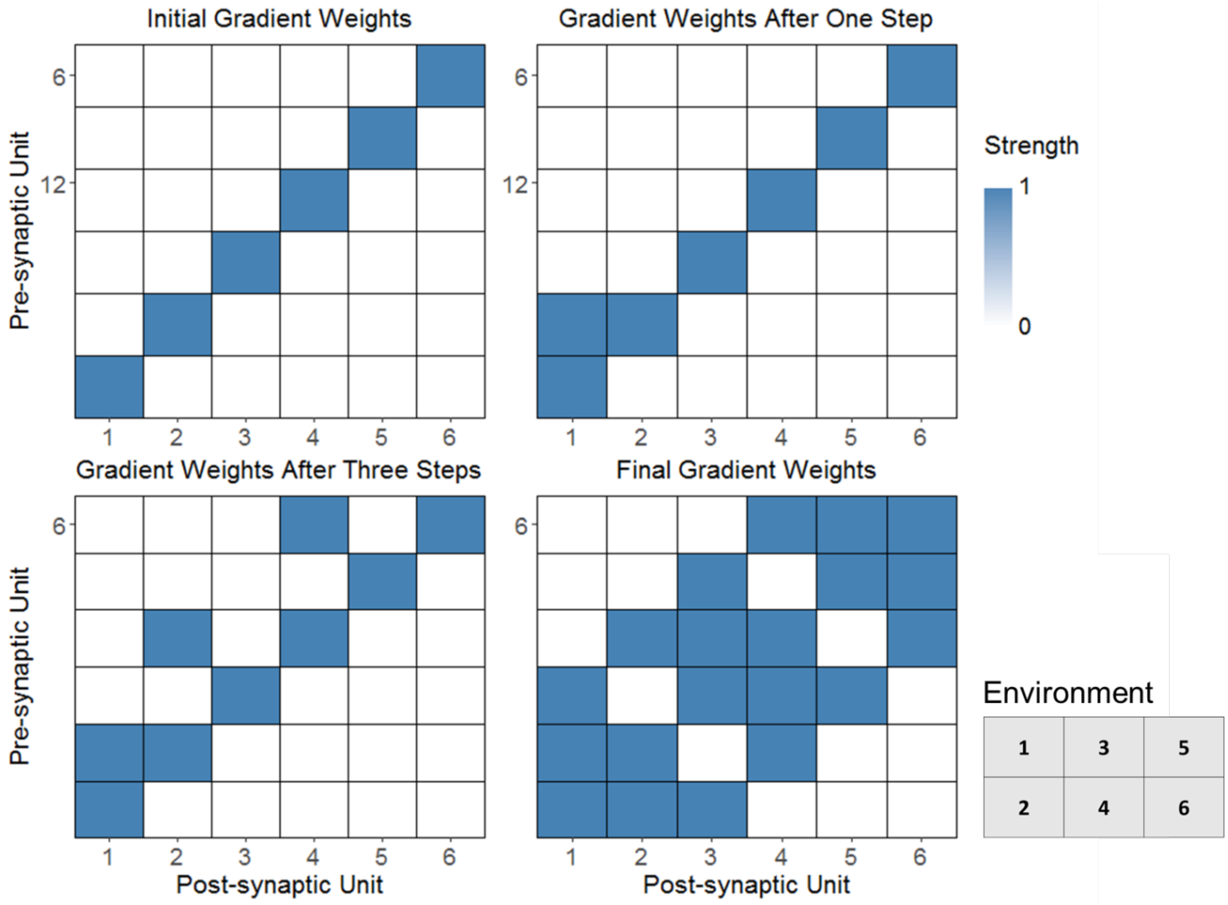


*Figure S9.* Progression of goal gradient weights starting with the initial values in the upper left, the weights after one step (from state 1 to state 2) in the upper right, the weights after a full trial in the lower left, and the final configuration of weights after all learning in the lower right. For example, state 6 can be reached by states 4, 5, and 6 so unit 6 receives input from units 4, 5, and 6. Note that units 3 and 4 receive input from more units than the others due to the fact that states 3 and 4 are adjacent to the most other states, which biases the gradient in their favor.

**Transition to Action**

The final projection requiring learning in the core model is the projection from the *transition-output* to *action-input*. During the acting phase, *transition-output* receives input from d*esired-transition* which in turn receives input from *current-state* and *next-state*. Each pair strongly activates a single *desired-transition* unit according to hard-coded weights implementing a simple conjunctive encoding. After learning, activation of a desired transition via *transition-output* should strongly excite the unit corresponding to the action which will effect that transition.

Learning the transition-to-action mapping again raises the dual roles problem. Increasing the strength between a transition and action only when the action successfully implemented the desired transition would be too inefficient, discarding information from all trials in which the action didn't have the intended consequences. Learning would only take place when the randomly chosen action happened to correspond with the desired transition, which is unlikely to occur if the number of transitions and possible action are large.

A more efficient solution is for learning to occur every time a transition is implemented, connecting the observed transition to the previously taken action (Figure S10). However, this requires maintenance of the previously taken action and the observed transition, whereas the transition and action layers typically represent the desired transition and the chosen action. As with the *goal-gradient* learning, the dual roles problem here can be solved using oscillatory gating of inputs. The *transition-output* layer must pull double duty, representing the desired transition in the pursuit of the goal, but representing the observed transition to facilitate the appropriate learning. Likewise, the *action-input* layer must represent the action that will most likely implement the desired transition in the pursuit of the goal, but represent the action responsible for the previous transition to facilitate the appropriate learning.

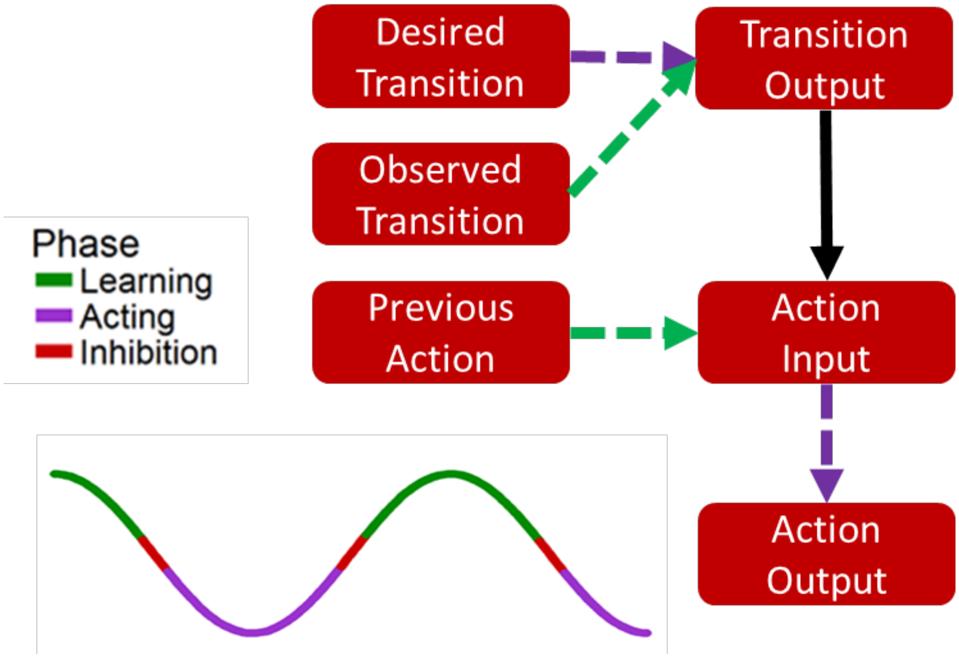


*Figure S10.*. Subnetwork responsible for learning the transition to action mapping. During the learning phase, transition-output and action-input get information about the previously observed transition and the action that caused that transition, resulting in Hebbian learning in the projection between the two layers. The projection to action-output is blocked during this phase to prevent the agent from repeating the pervious action. During the acting phase, a desired transition drives the associated action.

Additional layers maintain the past experience of the agent to be used during the learning phase. Information about the previous state of the network is maintained across two layers, *previous-state-1,* and *previous-state-2*. As shown below, the *previous-state-1* layer receives input from state but has a high time constant ($\tau$ = 4), which causes activity to build and decay slowly (Figure S11). *Previous-state-2* performs winner-take-all (WTA) on the input from *previous-state-1*. The inhibition at each state transition restarts the competition and since the activity corresponding to the new state has not yet caught up to the slowly decaying activity representing the old state in *previous-state-2,* the unit corresponding to the previous state becomes dominant. Together, the previous state and current state define an observed state transition, just as the current state and desired next state specify a desired transition.

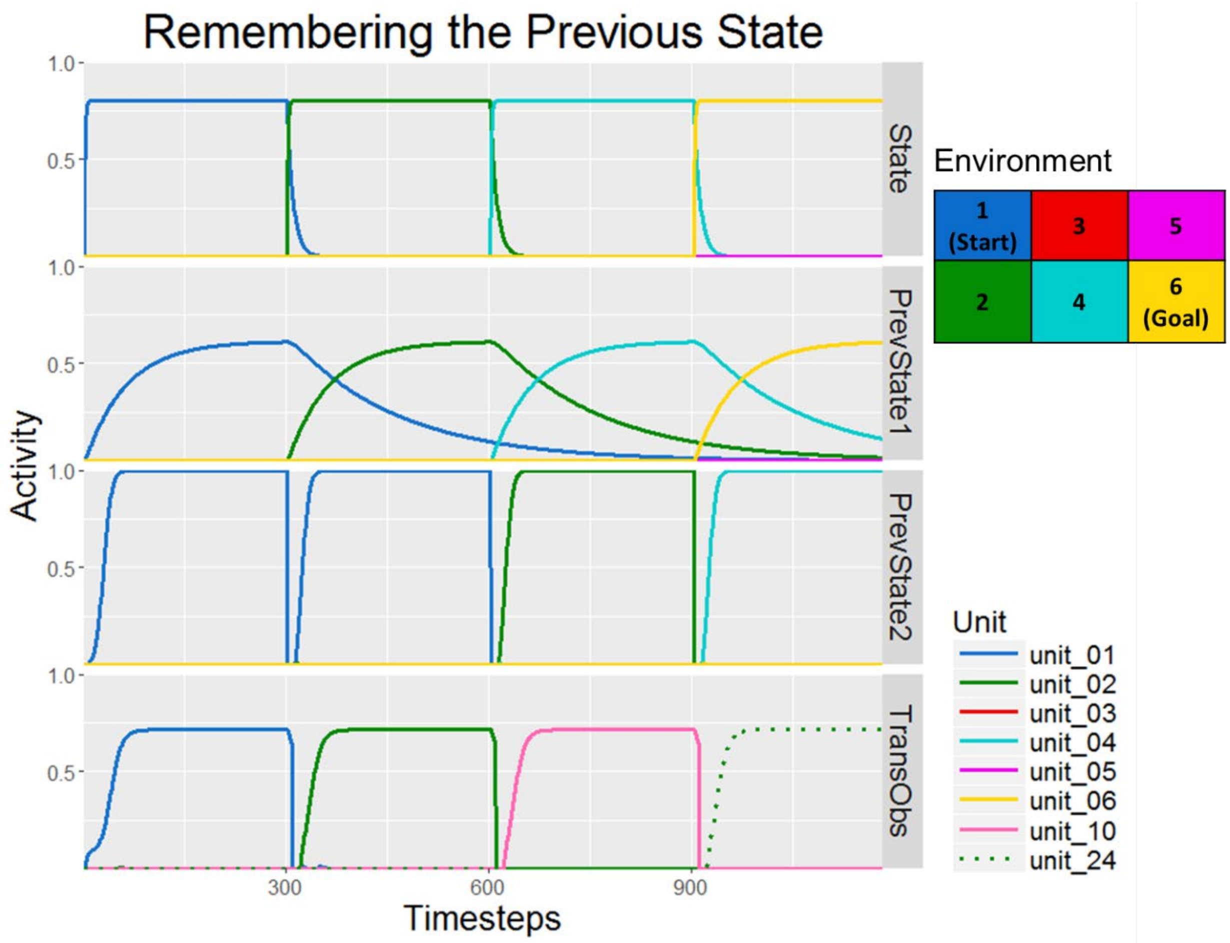


*Figure S11.* Model activity in layers responsible for monitoring the past state transitions during one trial. The first three layers have six units, one for each state. The observed-transition layer (abbreviated TransObs) has 36 units, one for each transition. After the first state transition, for example, the current state is state 2, the previous state is state 1, and the observed transition is transition 2. The final observed transition is transition 24, from state 4 to state 6.

The previous action is represented in a simpler fashion, with a layer, *previous-action,* that receives input from the *body* node and maintains it until the next state transition, as shown in Figure S12 below.

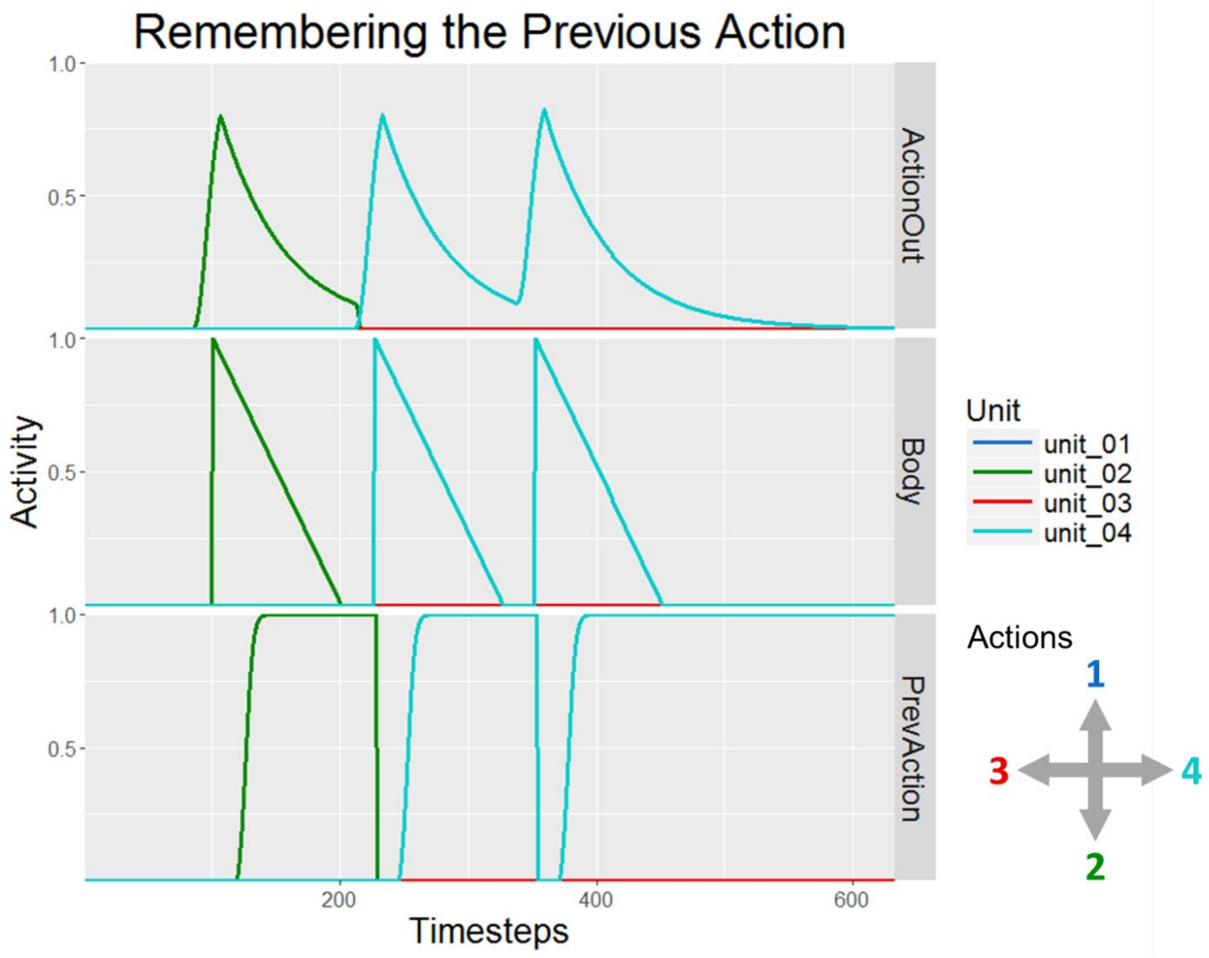


*Figure S12.* Model activities in layers responsible for monitoring the previous actions. Each layer has four units, representing movement in the four cardinal directions. *Body* is a node that registers above-threshold activity from action-output with a spike in value to 1. The linear decay of the signal enforces a minimum time period before the next action can be taken (a new signal won't be triggered in body until it has decayed to 0). Previous-action keeps a record of the last observed action until a new state transition occurs.

During the learning phase, t*ransition-output* and *action-input* receive dominating input from *observed-transition* and *previous-action*, and learning turns on shortly after these projections are opened. At the same time, the projections from *desired-transition* to *transition-output* and from *action-input* to *action*-output are blocked. The weights between *transition-output* and *action-input* are updated according to,

$$\dot{w} = \theta\eta\left[\left(\left(a_{pre} > \varphi\right) * \left(a_{post} > \chi\right)\right) - w\right]^{+}, \tag{S8}$$

which is similar to the learning laws already seen, though learning is based on both pre and post synaptic activity rather than any eligibility traces. In this projection, $\varphi = .42$ and $\chi = .15$. These thresholds are chosen to be slightly below peak activity for their respective inputs. Using this learning law, these weights evolved over learning as shown below in Figure S13.

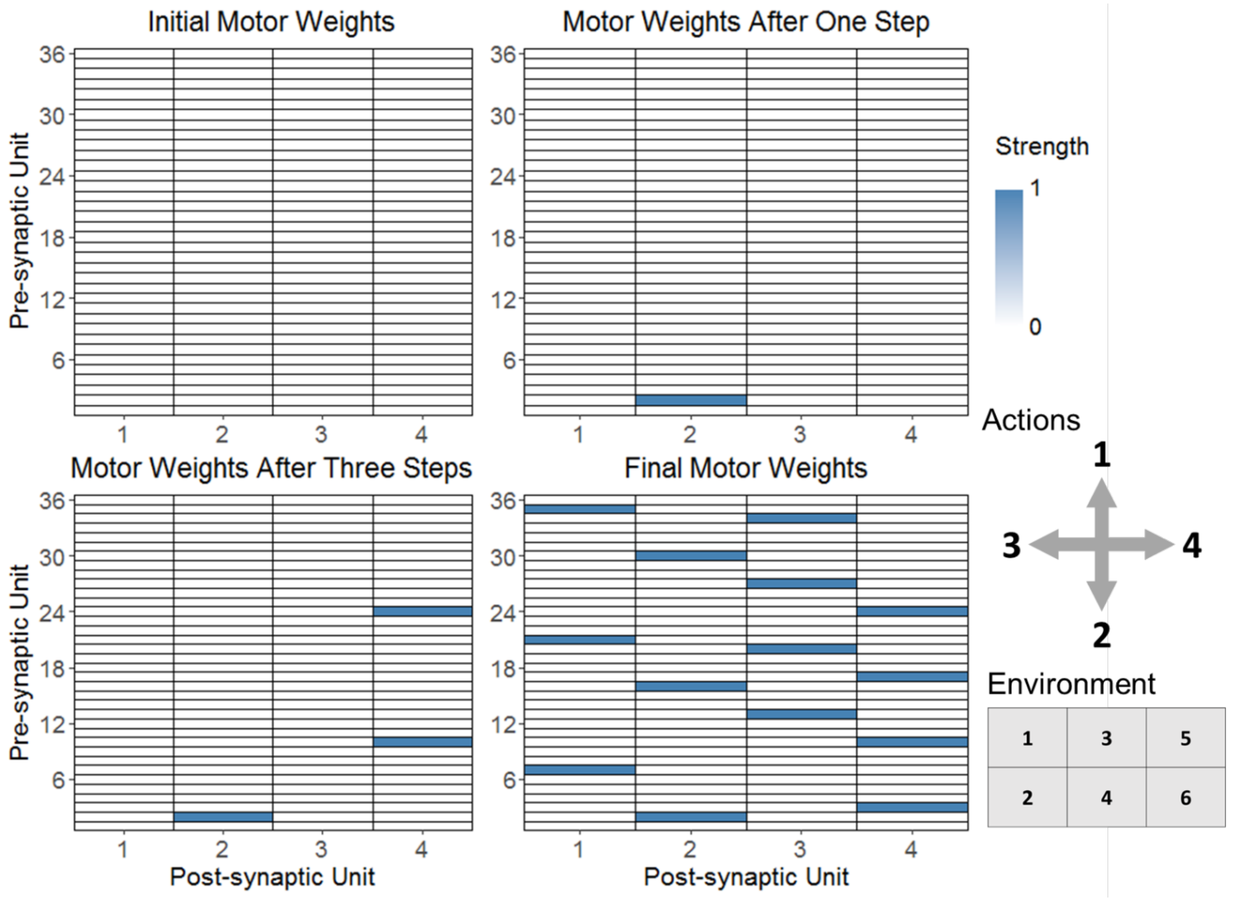


*Figure S13.* Progression of weights during learning in the projection from transition-output to action-input. Initialized at zero, the weights increase toward one when an action is observed to have resulted in a state transition. The six possible transitions from state 1 are numbered 1-6, while the possible transitions from state 2 are numbered 7 – 12, etc.

During the "acting phase" of the *substitution-oscillation*, *transition-output* reflects *desired-transition* and the activity in *action-input* is determined by the learned weights in the projection between *transition-output* and *action-input* while *action input* passes its activity pattern to *action output* in order to drive behavior.

**Extra Modules**

In this section we describe some additional model components. These are not strictly necessary for the core model to perform its function of learning goal-directed action, but the additional components provide abilities to (1) learn which goals should be activated given basic drive representations, and (2) plan a sequence of actions, and store that sequence, prior to execution.

*Goal Learning*

The behavior of the model is driven by activity in *goal*, in that the agent uses the goal signal as a set point for the current state signal and attempts to bring the current state into alignment with the goal. In the simulations described in the main text, *goal* activity was imposed externally via a node. However, goals can be learned and established via exploration with the inclusion of a few additional model components. A major motivation for creating a model that could pursue any goal state rather than traditional “reward” is that the value of environmental states can vary with the agent’s needs and desires. In goal-learning simulations, the agent’s needs are represented in a layer *drives* which, due to input from a corresponding node, has units whose activity increases slowly over time, like hunger or thirst. Each drive can only be reduced by reaching particular states of the environment (Skinner & Hull, 2006). If at agent reaches the appropriate state then, after a brief delay, the drive corresponding to the discovered reward is reset to 0.

The new *drives* layer projects to *goal* via a learned projection (Figure S14). The learning target is for each unit of *drives* to strongly excite the *goal* unit corresponding to the state(s) in which the drive is satisfied. For example, if state 6 contains a particular type of reward, the drive associated with that reward should activate state 6 as a goal. To learn the appropriate mapping, the projection must have information about both the presence of the reward and as well as the current state. The presynaptic component of the learning law is therefore a vector that indicates which drives have recently decreased at a rate below some threshold. The postsynaptic component cannot be activity from the *goal* layer, since the activity in *goal* represents the target state rather than the current state. Therefore, the projection receives modulatory, one-to-one, input from *current-state*[2]. This input does not activate the corresponding units in *goal* but does determine which units' weights are eligible to learn. If the agent is in state 2, the weights on unit 2 in *goal* are eligible to be modified and will increase when a drive is substantially reduced while in that state. The learning law governing this process is,

$$\dot{w} = \eta\Big[\Big((\dot{a}_{pre} < \varphi) * (a_{mod} > \chi)\Big) - w\Big]^{+} \qquad \text{(S9)}$$

The presynaptic component of the learning law requires the relevant drive to be currently dropping at a rate less (faster) than $\varphi$ and the postsynaptic component requires that the modulatory input from *state* be above a threshold $\chi$. In short, if the agent is in a state where a drive is decreasing, the agent will learn to return to that state the next time that drive is high.

[2] It would also have been possible to use an oscillatory solution with *goal* receiving input from *drives* and *state* in acting and learning phases, similar to the process used in other projections.

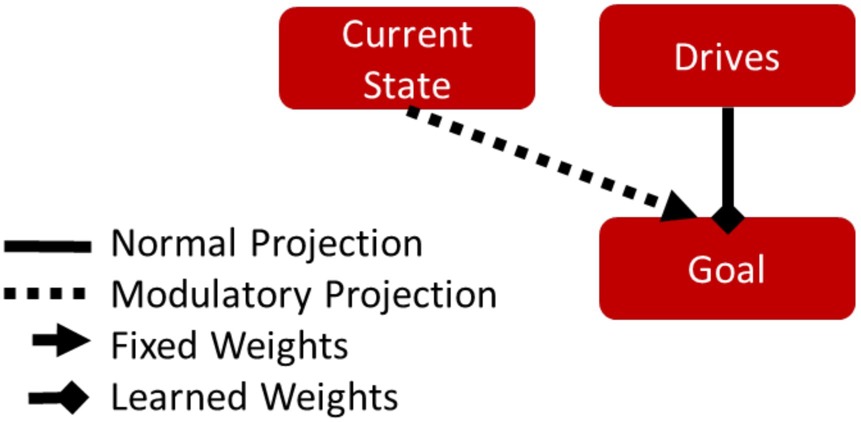


*Figure S14.* Layers involved in learning which goals to pursue based on drives.

As described, this scheme will work as long as only one drive is high. However, if two drives become highly active, their respective goals may interfere with each other. To address this, winner-take-all dynamics within *goal* help stabilize the activity pattern. To allow goal changes reflecting changing motivational drives, inhibition associated with state changes was added to *goal*, similar to the inhibitory pattern in many other layers. If, after a state change, the relative strength of two drives has changed, the selected goal will shift accordingly.

After learning the drives-to-goal mapping in addition to the transition structure and action mapping, the agent can autonomously navigate an environment in order to satisfy its changing needs. To demonstrate this, we put the agent in an environment with the following configuration.

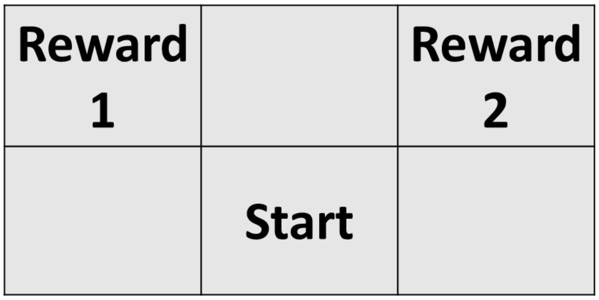


*Figure S151. Small grid-world environment with distinct rewards in different states. Based on the levels of two drives, the agent must decide whether to pursue reward 1 or reward 2.*

After only one exploratory trial, the network had learned which states contained the reward in addition to the layout of the state-space and the mapping between actions and state transitions. On a subsequent trial, illustrated below in Figure S16, both drives were initially active. The agent first adopted the goal of reaching state 5, where reward 2 was present. After successfully reaching this state, the associated drive diminished and the agent successfully navigated to state 1 to satisfy the remaining drive. On a following trial, shown in Figure S17, the drives were not activated initially but left to increase gradually over time. The agent has no goal and moves around the environment randomly until the first drive becomes sufficiently active and goal 1 is instantiated. The agent then quickly navigates to state 1, reducing the active drive.

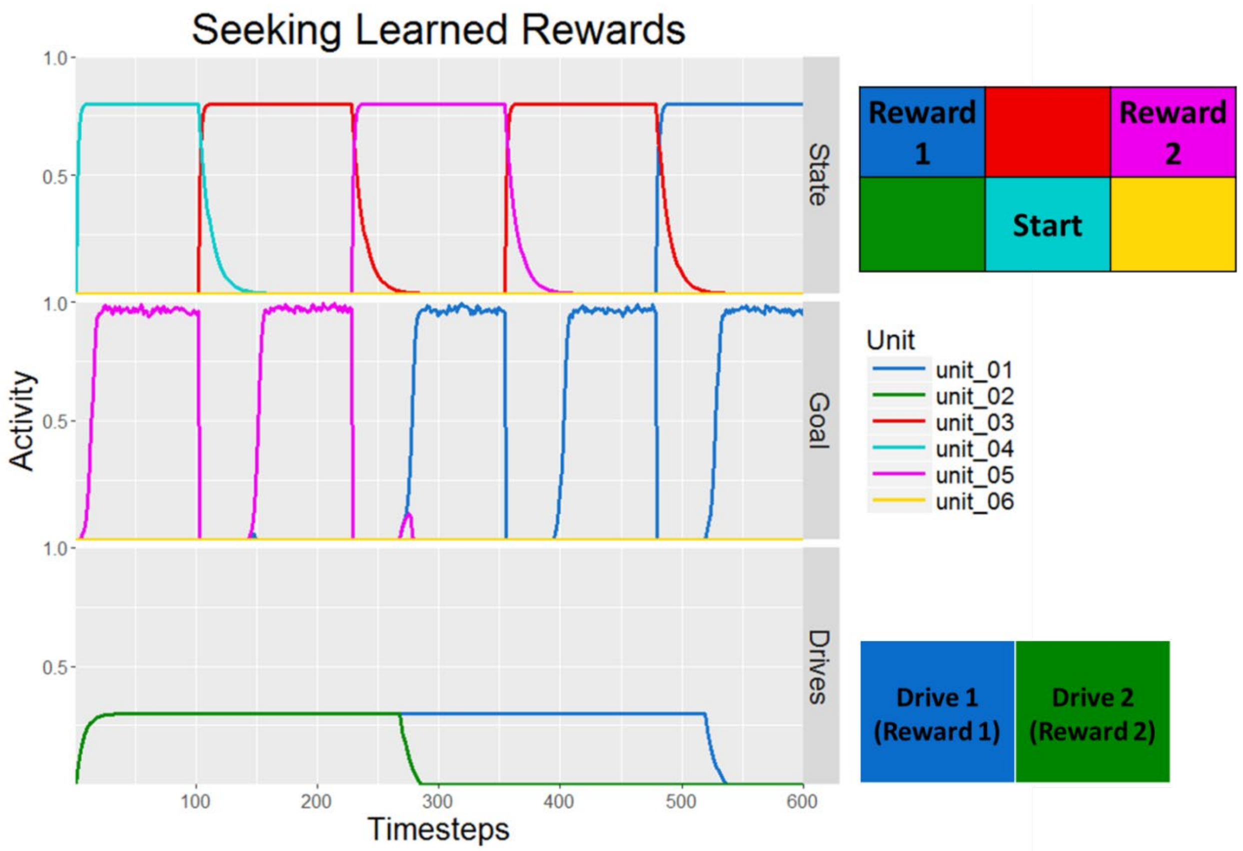


*Figure S16.* Network activity in layers related to goal selection based on drives. The current-state and goal layers have 6 units, one for each state, while drives has two units corresponding to the two possible rewards. Early in the trial, both drives become equally active and the network choses to pursue reward 2 in state 5, which it quickly reaches. The drive associated with that reward drops to 0 and the network then selects state 1, the location of reward 1 as the goal state and quickly navigates there.

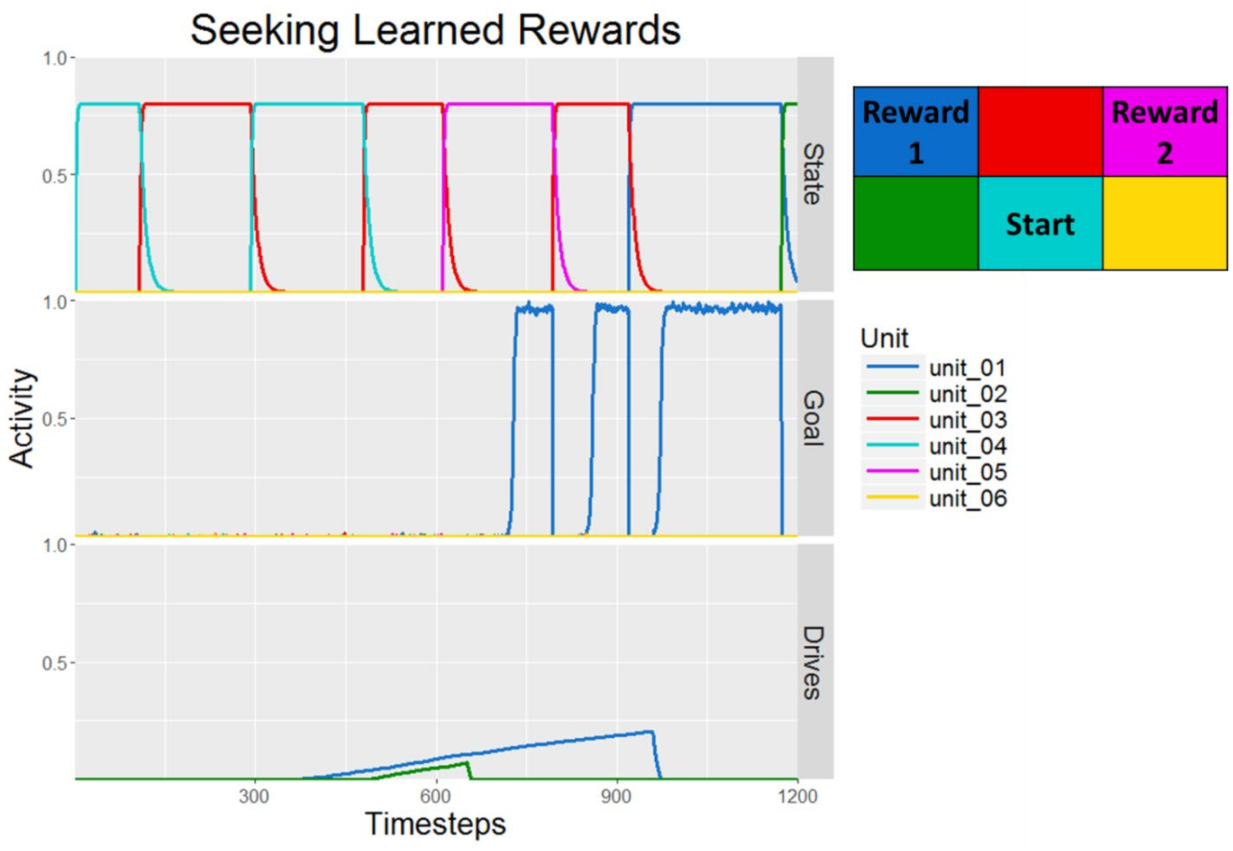


*Figure S172.* Network activity in layers related to goal selection based on drives. In this trial, the value of drives increases semi-randomly. Once drive 1 is sufficiently active, state 1 is established as a goal state and the network quickly moves there to reduce the drive values back to baseline.

### Planning

Because the *goal proximity* gradient represents the adjacency of each state to a potentially distant goal, the GOLSA agent is not *myopic* in the typical RL sense referring to an agent with only information about its immediate surroundings (Sutton & Barto, 1998). However, at each step the agent only "plans" one transition ahead, even though in many real-world cases it is useful for agents to store multi-step plans in advance of execution. The ability to store and carry out an action plan can be added to the GOLSA model with a module consisting of several components and connections.

The core component is a simple competitive queue where the activity level represents temporal priority (Daniel Bullock & Rhodes, 2003). Units in the main queue layer, named *queue-store*, maintain the planned trajectory using units with a decay parameter of 0, allowing excitation levels to stably persist. The degree of activity codes for temporal precedence, such that the movement encoded by the most active unit is executed first. Actions, conceived of in the spatial navigation case as “move left,” “move right,” etc., cannot be adequately stored this way as there is no ability to represent action repetition. If, during sequence storage, the unit in *queue-store* representing “go left” is excited twice in order to represent two moves to the left, its activity will increase above the level of other units which were activated only once and the network will interpret that activity as indicating “go left” should be executed soon, rather than twice. Therefore, the queue stores sequences of state transitions rather than actions per se. The queue layers, like the transition layers, therefore contain $n^2$ units where n is the total number of possible states.

This scheme requires a number of modifications to the existing network as shown in Figure S18. First, a control node, *queue-mode*, is used to indicate to the network whether the queue is in loading mode, execution mode, or inactive. A new layer, *simulated-state*, represents the state the network “imagines” itself to be in during each step of planning. The queue itself involves three layers, *queue-input*, *queue-store*, and *queue-output*. When inactive, the projections connecting the queue to the rest of the network are gated closed (Figure S19).

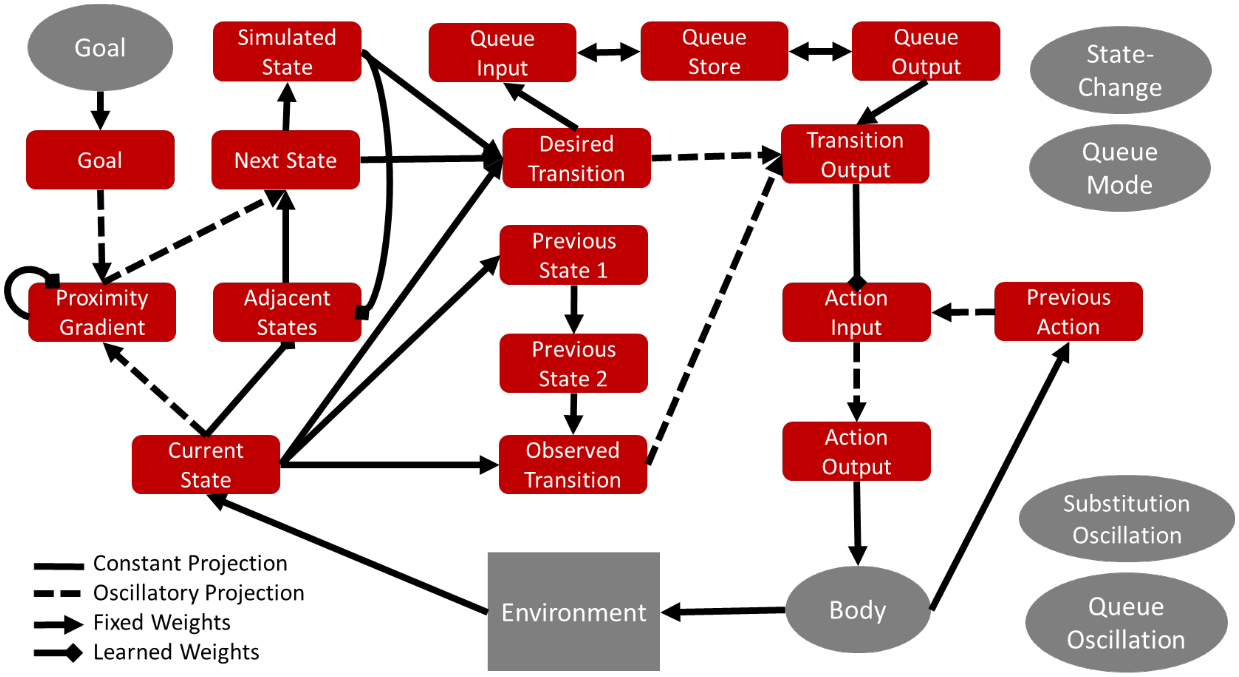


*Figure S18.* Network with queue module added (top row). Some node connections not shown. Also not shown is a one-to-one connection from current-state to simulated state required for learning the simulated-state to adjacent-states weights.

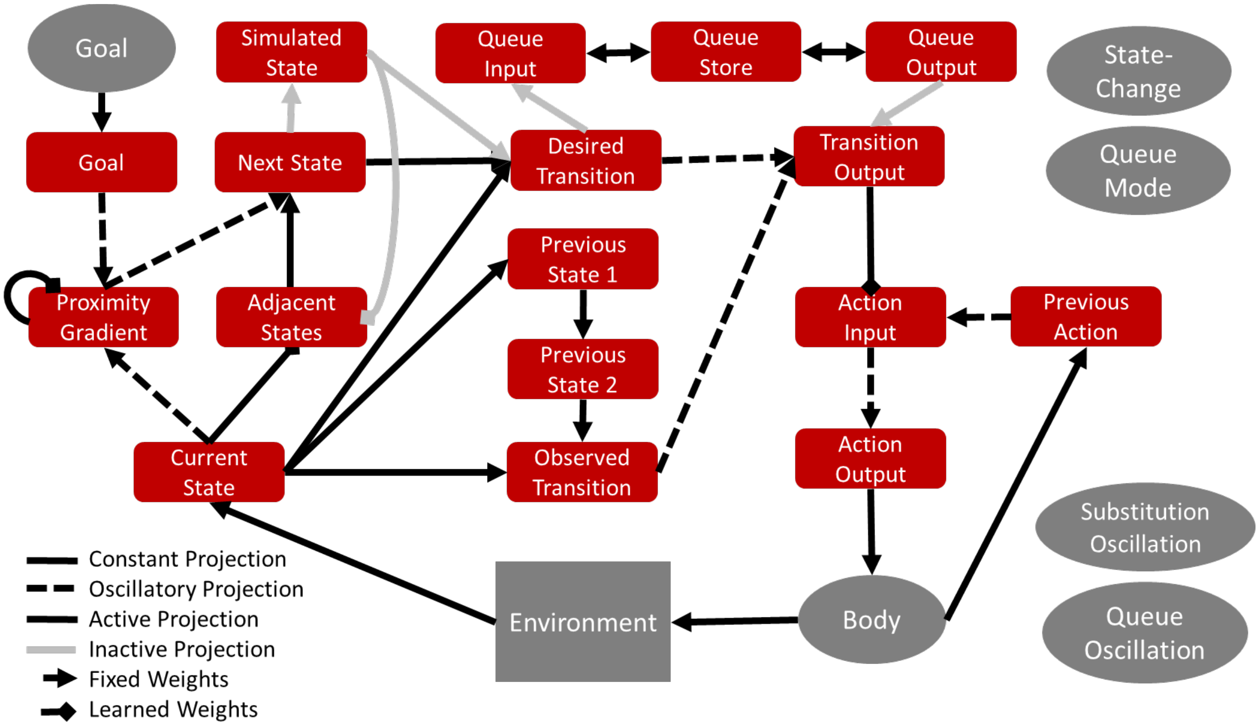

*Figure S19.* Network with queue module during non-planning mode. Projections to and from the queue module are gated closed.

#### Storing a Plan

While a plan is being stored (indicated to the network by the value of *queue-mode*), the network connectivity shifts to the pattern shown in Figure S20. In this configuration, *simulated-state* input replaces *state* input to *adjacent-states* and *desired-transition*, while all input to *transition-output* is blocked. This prevents actual action being taken while the network runs through a series of simulated transitions to reach the desired goal.

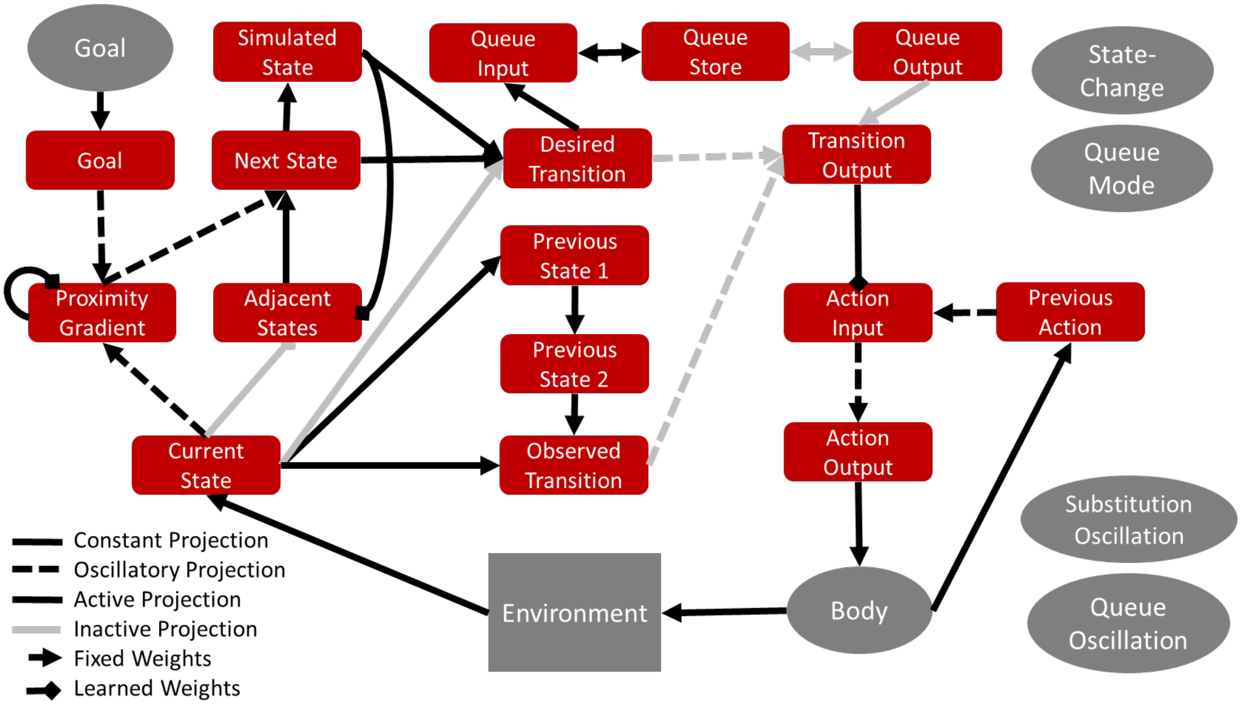


*Figure S20.* Network configuration with queue module in planning mode. Output to the motor system is blocked, simulated-state is substituted for state input, and desired-transition activity is fed into the queue.

The process of loading the queue with the desired state transitions is cyclical. It requires the gating and inhibition of various projections and layers in a particular order. Because this process is independent of the learning governed by the *substitution-oscillation,* a distinct oscillatory node *queue-oscillation* governs this gating pattern. The sequence can be summarized in three steps, illustrated in Figure S21, with the oscillatory mechanism providing a kind of clock cycle (Klausberger & Somogyi, 2008), as shown in Figure S22.

1. Replace the *simulated-state* pattern with the pattern in *desired-next-state* (simulating a movement to the desired next state).
2. The new *simulated-state* pattern will update the *adjacent-states* layer, resulting in a new state being selected in *desired-next-state*. Use the simulated state and desired next state to define a desired transition.
3. Load the desired transition into *queue-store* via a short burst of activity in *queue-input*.

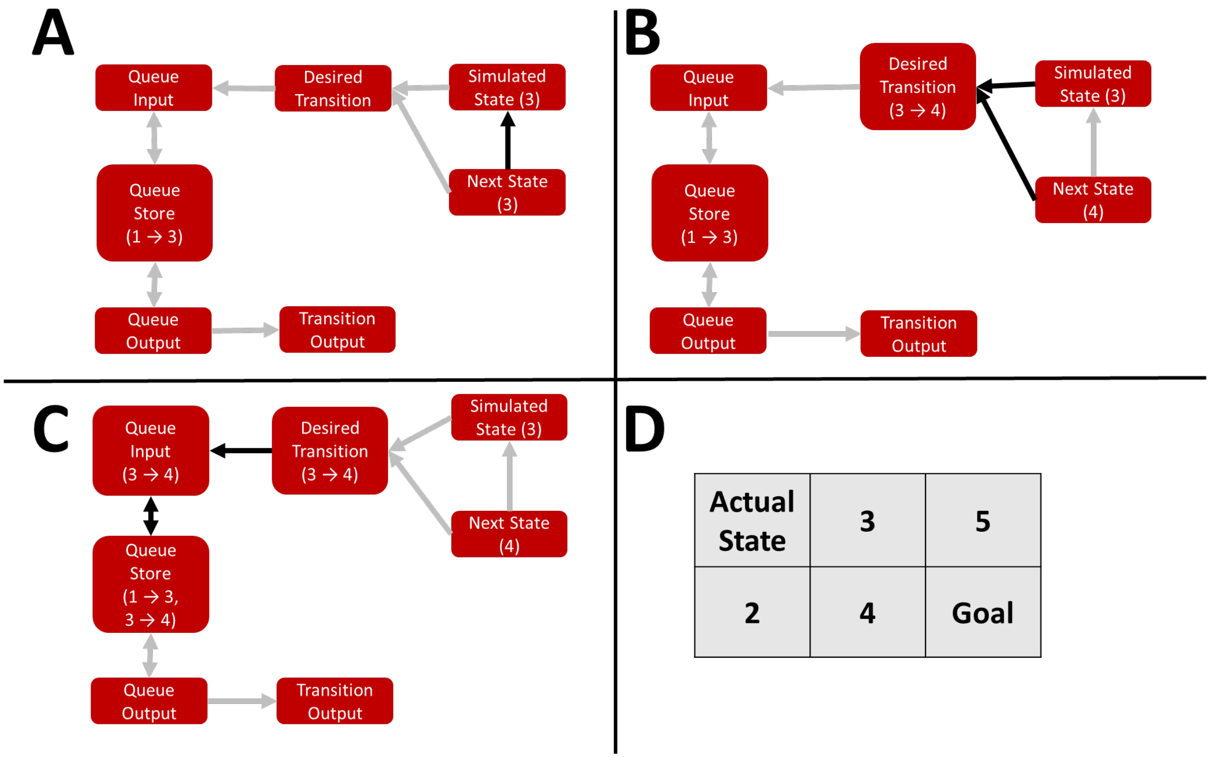


*Figure S21.* Illustration of the queue-loading process. The numbers or sequences in parentheses under the layer names represent hypothetical representational content of the layer. In step 1 (panel A), the simulated-state pattern is replaced by the pattern in next-state, representing an "imagined" transition between states. Panel B illustrates the second step in which *simulated-state* and *desired-next-state* (representing the best next state from the currently simulated state) specify a desired transition. In panel C, The desired-transition is gated into the *queue-store* for maintenance. Since a transition (from 1 to 3) was already being stored in the queue, inhibition from *queue-store* dampens the excitation in *queue-input* resulting in a less active trace for the second transition in *queue-store.*

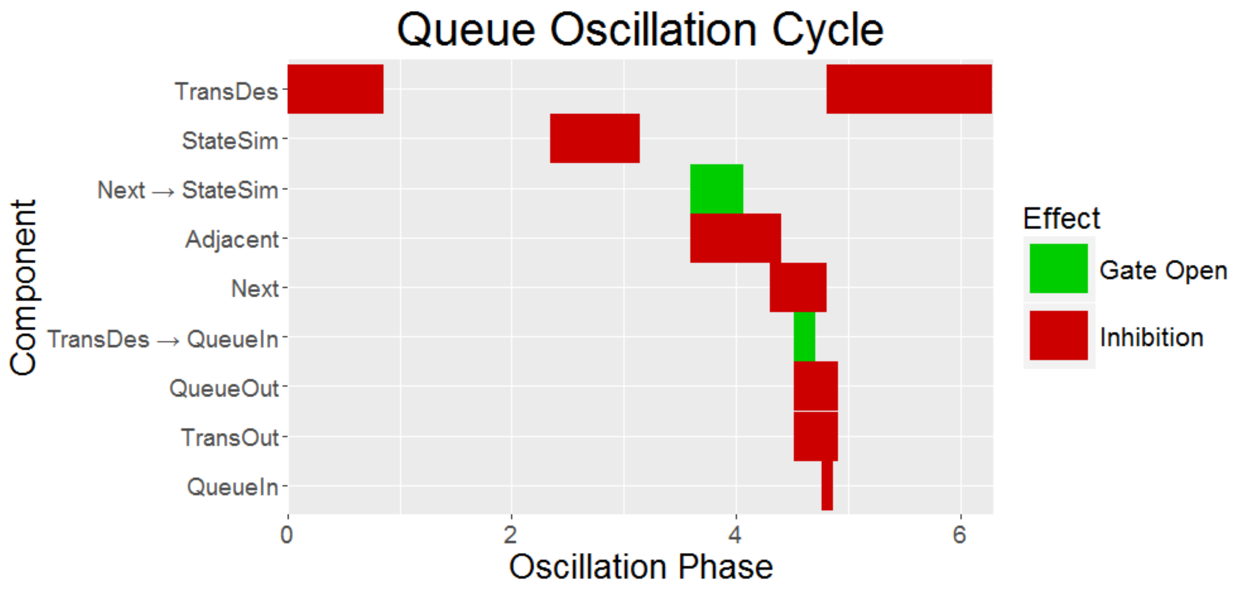


*Figure S22.* Sequence of inhibition and gating governing the queue-loading cycle. Starting with simulated-state, inhibition flows forward through the network, to adjacent-states, next-desired state, the queue layers, and then to *desired-transition* which remains inhibited until the next cycle. Within each cycle, brief windows allow input from *next* to *simulated-state* and from *desired-transition* to *queue-input*.

*Queue-in* receives very brief input from *desired transition* and relays that activity one-to-one to *queue-store*, which has a decay rate of zero and therefore keeps a continuous record of its inputs until actively inhibited. *Queue-store* exhibits a hard-wired, inhibitory all-to-all connection with *queue-input* such that the more units are active in *queue-store*, the more *queue-input* is inhibited and the less active the next input to *queue-store* from *queue-input* will be. This connectivity pattern encodes temporal precedence into the queue-in terms of higher stored activity.

The patterns exhibited by many layers in the network depend crucially on the patterns in upstream layers. As those upstream layers are updated, for instance after an imagined state transition, a burst of inhibition resets downstream layers allowing them to respond to the new input appropriately. The inhibition of each layer, along with the gating of key projections, is tied to the *queue-oscillation* such that inhibition or gating occurs at particular phases in the cycle, as shown above in Figure S22.

The entire queue-loading process can be observed below in Figure S23 which shows the queue-loading phase of a trial in the 6-state environment.

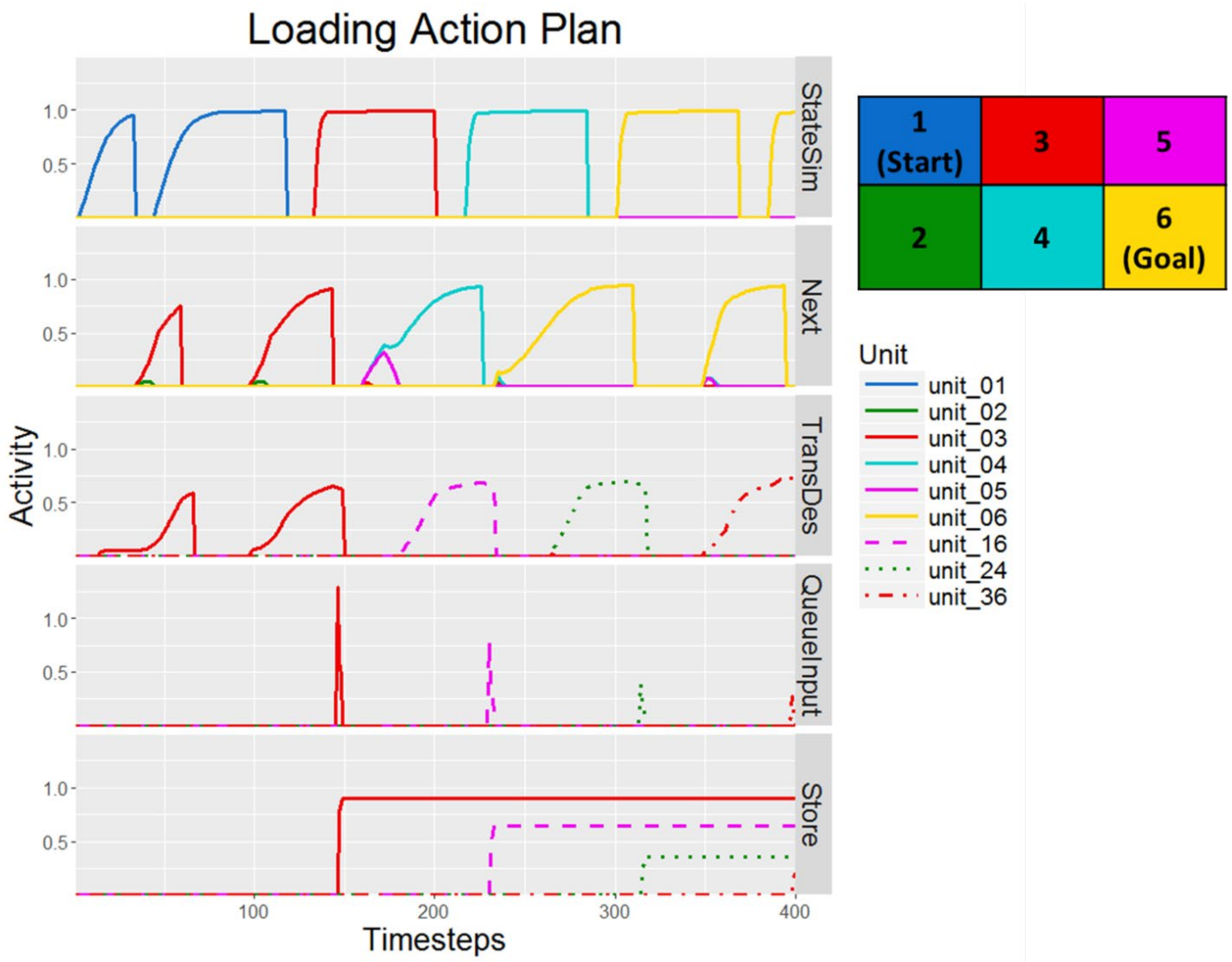


*Figure S23.* Model activities in layers related to queue-loading during one trial. At each step, the model selects a desired transition based on the activity in simulated-state and desired-next-state. This desired transition is briefly gated into queue-input which establishes a persistent trace in queue-store which codes for temporal order with activation strength.

As the *simulated-state* layer works through a trajectory from state 1 to state 6, *next-desired-state* represents the ideal next step from the current simulated state rather than the actual state (the agent remains stationary in state 1 during this period). The combination of values in *simulated-state* and *next-desired-state* activate the corresponding state transition in *desired-transition* which is briefly gated into the queue via *queue-input*. The resultant activation is maintained without decay in *queue-store* which displays a stable pattern of activity in which level of activation encodes temporal precedence. The most active transition unit represents a transition from state 1 to state 3, the next most active a transition from state 3 to state 4, and finally a transition from state 4 to state 6, the goal.

**Executing a Plan**

Network connectivity once again shifts when in plan-execution configuration (Figure S24). *Transition-output* receives input solely from *queue-output* while most of the rest of the network ceases normal operation, since the planning has already been accomplished.

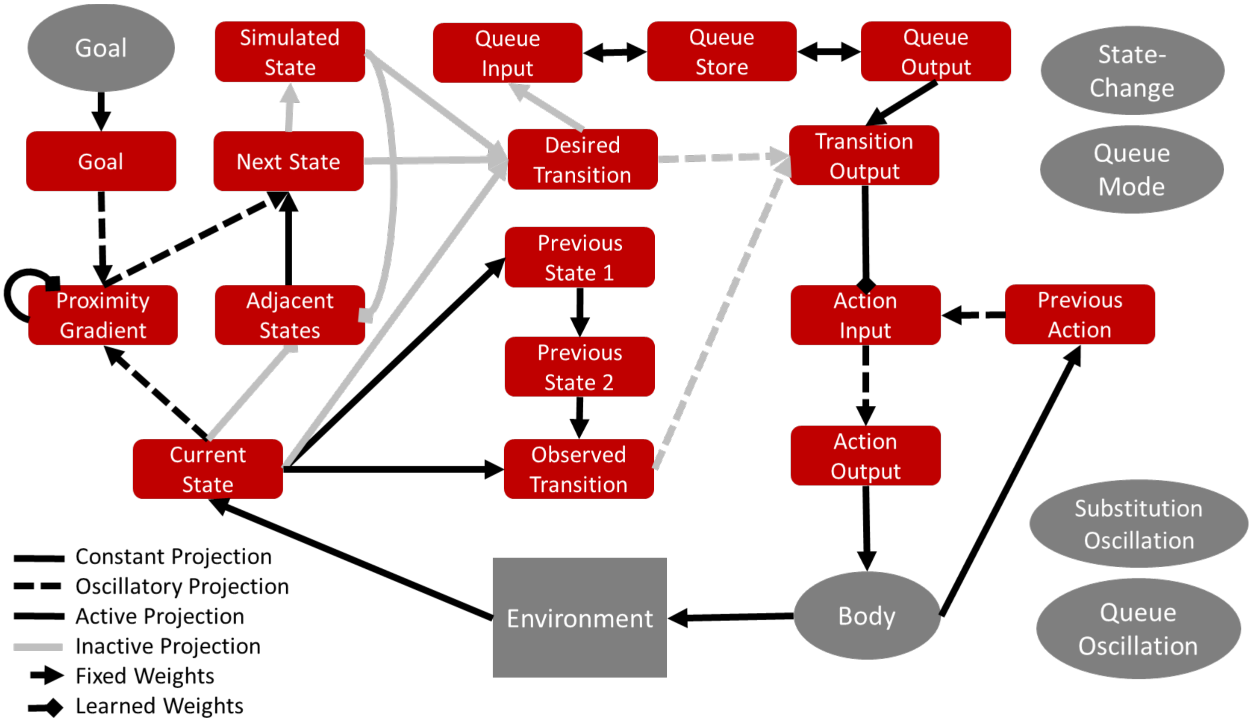


*Figure S24.* Network configuration in queue-execution mode. Most of the output from transition-selection layers is suppressed while the queue-store and queue-output layers drive action output that implements the stored transition plan.

Reading out a transition sequence relies on a similar cycle of activity to loading a plan, as depicted in Figure S25 below. As with queue loading, a pattern of inhibition and gating occurs continuously throughout the *queue-oscillation,* but the process can be approximately described as discrete steps (Bullock, 2004).

1. Gate open the projection from *queue-store* to *queue-out*. The WTA dynamic in *queue-out* results in activity only in the unit corresponding to the most highly active unit in *queue-store*.
2. After the competition has settled, send the desired transition to *transition-output* for access to the action system.

3. Delete the recently-executed transition from the queue by gating open the one-to-one inhibitory projection between *queue-output* and *queue-store*. *Queue-output* is then inhibited to clear it for the next cycle.

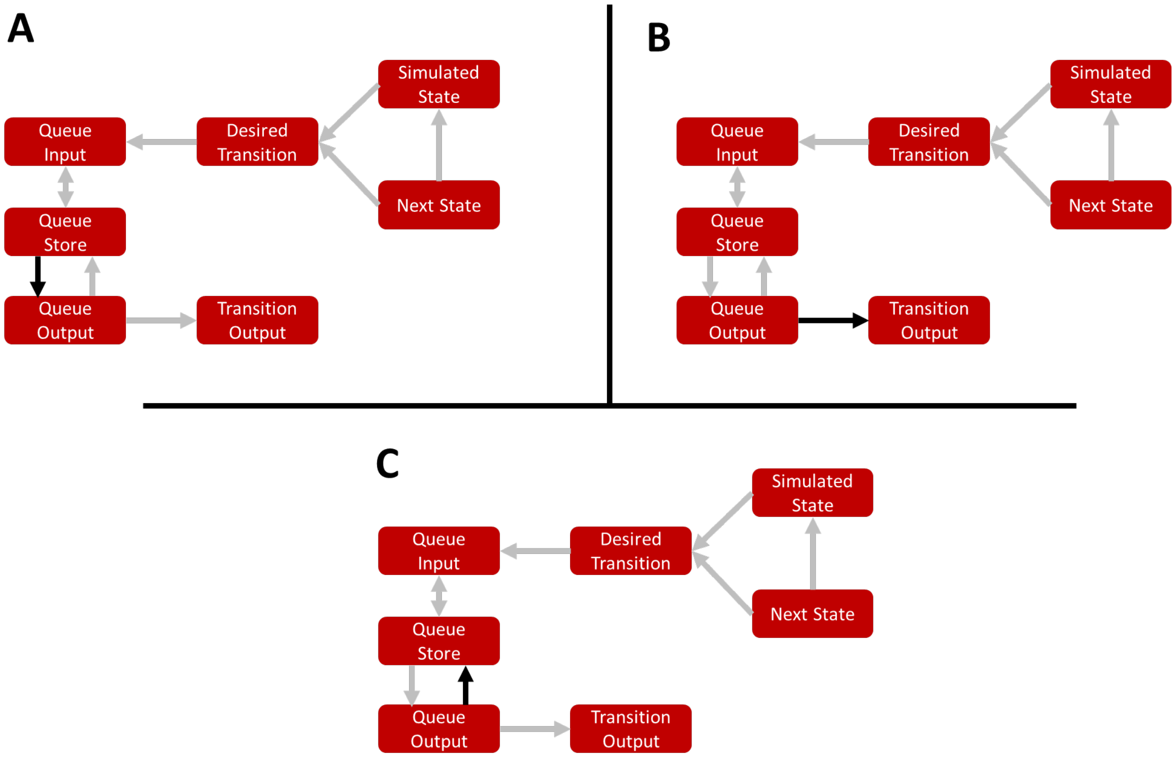


*Figure S25.* Illustration of the queue-execution process. First (panel A), queue-store is gated into queue-output. A WTA competition results in a single unit becoming active in queue-output, namely the unit corresponding to the most active unit in queue-store. The output is then gated to transition-output which drives action via a learned projection. Finally, the entry is deleted from the queue via an inhibitory projection from queue-output to queue-store.

Both the loading and the execution phases of a trial can be seen in the simulation results in Figure S26 below.

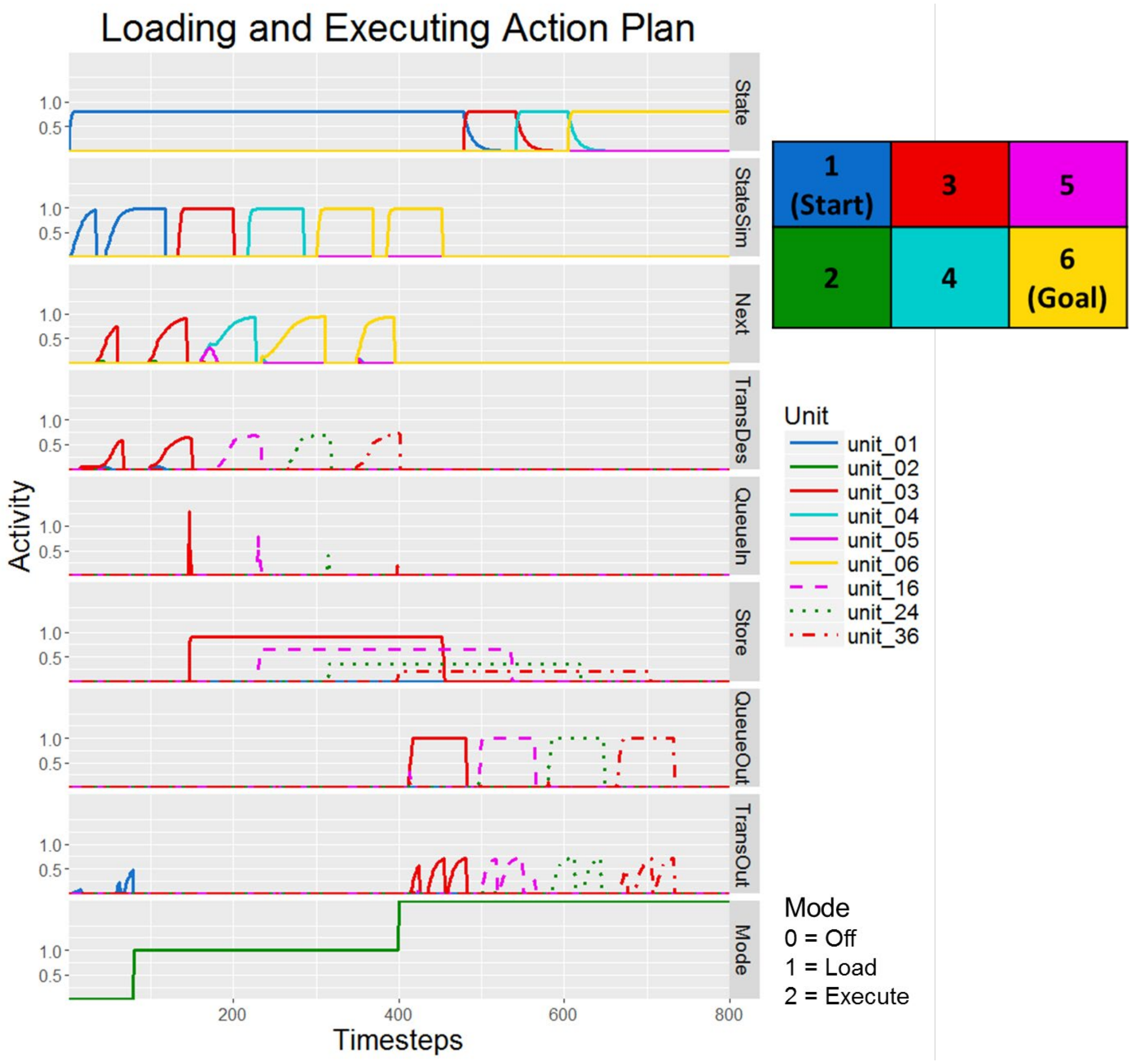


*Figure S26.* Network activities in layers relevant to queue loading and execution during a full trial (Figure 33 shows the first half of the same trial). During the queue-loading process (where Mode = 1) the agent remains in state 1 but simulates a sequence of state transitions to the goal, which are stored in queue-store. During the queue execution process, that sequence of transitions is gated into queue-output and then into transition-output which drives the appropriate transitions.

The data shown to the 400$^{th}$ timestep are identical to that shown in Figure S23, illustrating the loading of the queue. At that time, the network shifts into execution-mode and begins sending the stored transitions in *queue-store* to *transition-output* where they activate the necessary action in accord with previous learning. Shortly after each transition from *queue-store* reaches *transition-output,* it is deleted from the queue. In the trial demonstrated here, the agent successfully planned a series of transitions while stationary, then executed them in sequence to reach the desired goal.

### Biological Basis of Model Mechanisms

The GOLSA model was primarily constructed at the computational and algorithmic levels of analysis, with the objective of creating a system capable of controlling an agent in the service of goal-directed action (Marr, 1982). Because the model was created based on a high-level performance target rather than directly from observed data, and because the architecture is specified at a more abstract level than biological implementation, one concern is that it may be too unconstrained. It may provide a solution to the problem of goal pursuit that would be impossible or improbable to implement in a brain. However, each of the key mechanisms in the model is grounded in biological precedent, and we have been careful to observe the constraints of localist representation and learning, as well as the constraints of real-time activation propagation.

#### Components

Layers in the model are connected by projections, which consist primarily of a matrix of weights representing synaptic strengths between pairs of presynaptic and postsynaptic layer units. Positive weights indicate excitatory connections while negative weights represent inhibitory connections. In some projections, such as those used to enforce WTA dynamics on the target layer, the weight matrix contains both positive and negative values. For example, the layer *next-desired-state* has a recurrent connection such that each unit excites itself while inhibiting all other units in the layer.

In addition to layers and projections, the model includes nodes which can provide input to layers as well as providing control signals necessary for gating and layer-wide inhibition. Because they are both crucial for model function and not modeled in terms of biologically-derived activation functions, one might worry that they significantly diminish the neural plausibility of the GOLSA model. However, these concerns are largely unfounded. Models necessarily capture only a limited and simplified portion of real complex systems. The primary use of nodes in the model is to provide external inputs to the model system. The *environment* and *body* nodes represent perceptual input and motoric output, respectively. The environment node provides a one-hot input vector (i.e., where one element is 1 and the rest are 0) representing the agent's current location in the environment to the *state* layer which forms a similar representation, albeit one that is modeled neurally with shunting excitation, decay, etc. Similarly, for most of the simulations presented, a node provides input to the *goal* layer. With the goal-learning extension to the model, *goal* instead receives input from the *drives* layer, while the *drives* layer receives sole input from a drives node. The *goal*, *drives*, and *environment* nodes can be seen as summarizing the output of motivational and perceptual neural systems that are not explicitly modeled. The outputs of these nodes are simple and could easily be supplied by neural components in a more extensive model. Similarly, the *state-change* node which detects a change in the *current-state* signal could be instantiated neurally, though given the questions the GOLSA model was created to answer, the added complexity would be of little benefit.

**Learning**

One of the key advantages of GOLSA over MBRL as well as other neural models is its ability to learn about the environment in a way that respects the biological constraint of the locality of information, such that synapses grow stronger or weaker based only on signals that could plausibly be present in their vicinity. It is therefore imperative that the learning laws governing synaptic plasticity in the model are biologically plausible.

Learning occurs in three places in the core model, namely in the projections from *current-state* to *adjacent-states*, from *goal-gradient* to *goal-gradient*, and from *transition-output* to *motor-input*. The first two rely on eligibility traces as the pre or post-synaptic components of the learning laws described in Equations (S6) and (S7).

Eligibility traces ramp up with neural activity but decay more slowly, providing a signal that the given unit was recently active without exciting or inhibiting connected units. Despite their utility and widespread use in a variety of computational models, the precise biological substrate of eligibility traces remains somewhat unclear (Singh & Sutton, 1996; Sutton & Barto, 2012). Potential candidates include intracellular calcium concentrations or monoamine receptors located at the synapse (Frey & Morris, 1997; He et al., 2015; Schultz, 1998). He et al. (2015) found evidence of the latter mechanism in facilitating Hebbian learning in cortex, showing how traces for both synaptic potentiation and depression balance each other out to produce stable learning. Alternatively, if units are treated as subpopulations of neurons, eligibility traces could also be produced by the spiking of excitatory interneurons that excite local pyramidal cells below their firing threshold (Basu et al., 2016; Dudman et al., 2007).

Eligibility traces are typically thought to reside in the postsynaptic cell, producing learning when a high trace level is coincident with presynaptic firing. This scheme is replicated in only one of the learned projections, namely within the *goal-gradient* layer. The *transition-output* to *action-input* does not use eligibility traces, while the *state* to *adjacent-states* projection uses presynaptic rather than postsynaptic eligibility traces. For example, if unit 1 in *state* is highly active, all synapses from *state* are eligible to learn in the near future. After a state change, if unit 2 in *state* is now highly active, unit 2 in *adjacent-states* will also be very active due to the initial one-to-one connectivity, exceeding the threshold required to learn. Because the synapses from unit 1 will still have a strong eligibility trace, the synapse between *state* unit 1 and *adjacent-states* unit 2 will increase. Although presynaptic eligibility traces have been utilized in phenomenological models of spike-timing-dependent-plasticity (STDP), they are a bit harder to interpret biologically (Senn & Pfister, 2014). The challenge is to allow learning at synapses receiving input from a previous source, but not potentiate all nearby synapses. Hypothetically, the physical instantiation of "pre-synaptic" traces could still reside post-synaptically if they were localized to particular synapses, though direct evidence for this is lacking at present.

**Drive to goal learning**

The final learned projection, between *drives* and *goals*, is the most unusual is that the presynaptic component consists of the time derivative of *drives* while the postsynaptic component comes from modulatory input from the *state* layer. Under this scheme, if a drive drops in a particular state, the weight between the unit representing that drive and the unit representing that state in the *goal* layer is increased. The modulatory postsynaptic component could be biologically instantiated in two ways. First, the modulatory effect of the *state* input could create a very short lived eligibility trace in the *goal* layer cells. Alternatively, the *state* input could facilitate learning directly using a process similar to input-timing dependent plasticity (ITDP). ITDP refers a process observed in hippocampal cells wherein a cell population modulates learning between two other populations (the pre and postsynaptic populations) by stimulating the post-synaptic cells below threshold at a specific time relative to pre-synaptic input (Basu et al., 2016; Dudman, Tsay, & Siegelbaum, 2007). To the best of our knowledge, no observed neural plasticity rules depend on decreasing activity in one of the populations. However, the *goals* learning scheme in the GOLSA model is not particularly far-fetched. Given the complexity of neurobiology that gives rise to the wide variety of plasticity rules that have been observed, a neuromodulator system that increases in strength when firing rate starts high and then decreases seems well within the realm of possibility. Alternatively, the *drives* component could be made more complex with additional layers such that some downstream layer, e.g. *drives-decreasing* has units which are active only when the corresponding *drives* unit is decreasing. In other words, the complexity could be exported from the synapses to the structure of layers in order to perform the necessary computation.

### Dijkstra's Algorithm and A*

Though the GOLSA model operates as a dynamical system, it can be thought of as implementing the following algorithm. At each state, compute the distance of each adjacent state to the goal, then take the action that would implement that state transition that minimizes distance to the goal. After each transition, update the weights specifying distances between states and state adjacency. Algorithms that are somewhat similar enjoy widespread use in computer science, though of course they were developed without any requirement that they be neurally instantiated.

Dijkstra's algorithm is foundational to a variety of path-finding algorithms (Dijkstra, 1959). Each node represents a state. Given a starting node and a goal node within a weighted network (where the edge weights represent path length), the algorithm starts with no knowledge of the distances between nodes and aims to find the shortest path between the starting node and goal node. It exhaustively explores all nodes neighboring the starting node, storing the distance of each node from the start. Then the algorithm selects each of the recently explored nodes beginning with the node closest to the starting node, examining the distances of each of the newly selected node's neighbors from the starting node and storing the shortest observed path length from each node to the starting node. In other words, Dijkstra's algorithm explores outward from the starting node, keeping track of all shortest paths from the starting node to the edges of explored territory. Once the goal node is closer than any unexplored node, no additional information needs to be collected and the algorithm can return the shortest path between the starting and goal nodes. Dijkstra's algorithm is guaranteed to find the shortest possible path, but suffers from the drawback that it explores outward in all directions from the starting node, with no influence of the rough goal location on exploration direction. For instance, if Dijkstra's algorithm were used to find the shortest road path between Indianapolis and Los Angeles, it would explore many paths to the East Coast even though we know *a priori* that the solution will not lie in that direction.

The A* algorithm addresses this problem by changing how nodes are chosen for exploration (Hart, Nilsson, & Raphael, 1968). In Dijkstra's algorithm, the unexplored node closest to the starting node is chosen. A* also considers how far each unexplored node is likely to be from the goal node, using a heuristic measure which is selected by the user, within some constraints. As a result, exploration proceeds roughly in the direction of the goal.

The GOLSA model bears some superficial similarity to these algorithms in that it moves along nodes of a state space closer and closer to the goal. In some ways, the exploratory process of Dijkstra's algorithm also mirrors the diffusion of activity outward from the goal which produces the proximity gradient. However, these similarities are mostly superficial. First, it is important to distinguish the learning process from agent navigation. Dijkstra's algorithm and A* explore an environment to find the optimal path. The GOLSA model first explores an environment and can then leverage that knowledge at a later time to either navigate to the goal via the quickest route or form a plan composed of desired state transitions.

The theoretical framing of the problem, as well as the exploratory process, are also quite different. Both Dijkstra's algorithm and A* occur in an environment where the path length between adjacent states is variable whereas the GOLSA model operates in units of state transitions – each adjacent state is exactly one transition away. In addition, the GOLSA agent can only explore states adjacent to the current state, in contrast to the path finding algorithms which explore new states based on their distance from the starting state and/or goal with no topological restrictions. An exhaustive search through the state space in GOLSA would therefore involve considerably more backtracking than occurs in the Dijkstra and A* algorithms. Finally, nothing in the GOLSA model is analogous to the heuristic measure used in A* to prioritize searching particular branches of the state space for an efficient path. The largest similarity between GOLSA and A* is that when the GOSLA agent is acting in the environment, it moves to the state that an A* process would explore, namely the adjacent state that is also closest to the goal. However, given the constraints that the state space is discrete and in units of state transitions, this pattern is the only logical one for goal-directed action.
